# Inhibiting the right dorsolateral prefrontal cortex selectively enhances unsupervised statistical learning

**DOI:** 10.1101/2025.08.08.669288

**Authors:** Orsolya Pesthy, Zsuzsanna Viktória Pesthy, Teodóra Vékony, Karolina Janacsek, Dániel Fabó, Dezso Nemeth

## Abstract

The brain must balance the automatic extraction of environmental regularities with top-down cognitive control, yet the causal neural mechanisms governing this interplay are debated. In particular, the hemispheric contributions of the dorsolateral prefrontal cortex (DLPFC) remain unresolved. Here, we applied inhibitory repetitive transcranial magnetic stimulation (rTMS) to the left, right, or bilateral DLPFC in 95 healthy adults during a probabilistic sequence learning task. We found that inhibiting the right and bilateral DLPFC significantly enhanced statistical learning compared to sham stimulation, while the left DLPFC stimulation group remained similar to sham. Contrary to the hypothesis that this competition is mediated by the suppression of existing knowledge, episodic memory performance was unaffected across all stimulation groups. These findings challenge a direct episodic gating mechanism and suggest the DLPFC’s influence on statistical learning is hemisphere-dependent. As an alternative, we propose that right DLPFC inhibition shifts cognitive processing toward a more exploratory information-sampling strategy, a view supported by our finding of significantly greater reaction time variability in the right and bilateral stimulation groups. Together, our results provide causal evidence for a right-lateralized suppressive influence of the DLPFC on unsupervised learning and suggest its modulatory role is linked to information processing style rather than direct competition with episodic memory systems.

## Introduction

The brain resembles an ecosystem, wherein multiple cognitive processes interact and compete for influence over behaviour^1,2^. Adaptation to novel environments requires neural plasticity, long attributed to supervised or reinforcement learning, optimized by model-based processes. However, mounting evidence suggests that neural plasticity can also emerge through passive, unrewarded exposure to structured input, highlighting a fundamental role for unsupervised learning, which is in turn optimized by model-free processes^3,4^. Building on the hypothesis that unsupervised and supervised mechanisms shape representations jointly, we examine a critical instance of this process in humans: statistical learning (the acquisition of probability-based regularities without feedback or reward), and its balance with model-based memory retrieval. According to the competition hypothesis, model-free (such as statistical learning) and model-based (such as executive functions or episodic retrieval) processes rely on overlapping neural resources and may therefore inhibit each other, with the dorsolateral prefrontal cortex (DLPFC) serving as a key hub in controlling their balance^1,5–9^. One proposed mechanism is that the DLPFC acts as a gate to long-term memory –-thus, its inhibition hinders retrieval while simultaneously supporting the encoding of novel statistical structure^9^. To test this hypothesis, we applied inhibitory repetitive transcranial magnetic stimulation (rTMS) to the left, right, or bilateral DLPFC in healthy participants, assessing its impact on both statistical learning and episodic memory retrieval.

The antagonistic relationship between DLPFC activity and the model-free process of statistical learning has been well-documented. Specifically, reduced DLPFC engagement has been consistently associated with enhanced statistical learning performance. This has been observed in various groups with reduced DLPFC activity, such as children (compared to adults)^10–12^, individuals under hypnosis^13^ or depletion^2,14,15^. Importantly, studies using inhibitory non-invasive brain stimulation on the bilateral DLPFC have shown better learning^9^, and retrieval^16^ of probabilistic information in the DLPFC-inhibited group compared to the controls. These findings together suggest an inverse relationship between DLPFC activity and statistical learning, demonstrated through both correlational and causal evidence (although see ref^17^) –-However, this alone does not clarify which mechanism drives the competition.

The hemisphere-dependent involvement of the DLPFC in statistical learning may offer valuable insight into its underlying mechanisms. Neuropsychological studies have shown a right hemisphere dominance in statistical learning, highlighting that statistical learning remains intact when solely the right hemisphere remains available in split-brain and unilaterally lesioned patients, whereas the left hemisphere supports systematic hypothesis-testing - essentially, model-based functioning^18,19^. This asymmetry seems to extend to the role of the DLPFC, where improved statistical learning has been observed following inhibition of the left or facilitation of the right DLPFC^14,15,17^. However, findings remain inconsistent, with some studies reporting the opposite pattern^20–22^ or null results^23,24^. A proposed mechanistic explanation is rooted in hemispheric differences in decision strategies: the right hemisphere favors selecting the most probable option each time (frequency-maximizing strategy), while the left hemisphere engages in choosing each option in proportion to its occurrence frequency (frequency-matching strategy)^19^. While the latter supports causal inference, statistical learning, in which frequency is the key information source, may benefit more from frequency-maximizing strategies^4^. Although this account is compelling, the lateralization of statistical learning remains unresolved due to inconsistencies in the literature. To move beyond descriptive accounts and clarify the role of DLPFC asymmetry, we applied repetitive transcranial magnetic stimulation (rTMS) over the left, right, and bilateral DLPFC, and examined its effects on both statistical learning and episodic memory retrieval.

In the domain of episodic retrieval, most findings align with the competition hypothesis and the DLPFC’s proposed gatekeeping role in regulating access to long-term memory. A positive relationship between DLPFC functioning and episodic retrieval performance has been shown, although null results are common as well (for a review, see ref^25^). This pattern suggests that the DLPFC may modulate memory access, possibly through a working memory involvement^26^. However, it remains unclear whether downregulating episodic retrieval facilitates the encoding of new statistical regularities^9^. To directly test this, we included an episodic memory task in our study, allowing us to examine whether DLPFC modulation affects retrieval performance and whether this, in turn, relates to statistical learning outcomes. Hemispheric asymmetries offer further insight. According to the classic Hemispheric Encoding/Retrieval Asymmetry (HERA) model^27,28^, the left prefrontal cortex plays a greater role in encoding, while the right is more engaged during retrieval. Additional factors, such as stimulus modality (verbal stimuli are more left-lateralized, visual stimuli more right-lateralized), emotional valence (with mixed findings), and familiarity (with the DLPFC particularly engaged in the recall of novel stimuli), further shape lateralization effects^25^. To control for these variables, our memory task used novel, emotionally neutral stimuli incorporating both verbal and visual elements. Crucially, if increased statistical learning is observed in tandem with impaired episodic retrieval –-especially following right DLPFC stimulation –-this would lend support to the hypothesis that the DLPFC acts as a cognitive gate to long-term memory. When this gate is weakened, the system may become more receptive to encoding novel statistical patterns.

We aimed to address open questions about the effects of DLPFC inhibition on statistical learning and episodic memory. We applied 1 Hz (inhibitory) rTMS over the left, right, or bilateral Brodmann 9 area to target the DLPFC. In this sham-controlled study, stimulation was on in the breaks of task performance when participants performed on a statistical learning and an episodic retrieval task in a randomized order. Furthermore, we re-tested the participants after a 24h delay. We had the following expectations regarding the intervention effect:

1. In this study, we go beyond the findings of Ambrus et al.^9^ by addressing a key open question: the hemispheric asymmetry in statistical learning. Specifically, we aim to identify which hemisphere plays a more critical role in this process. If 1 Hz inhibitory rTMS over the left hemisphere enhances statistical learning performance, it would support right-hemisphere dominance in statistical learning, emphasizing the role of different cognitive strategies as a background mechanism^19^. Conversely, if altered excitability of the right dorsolateral prefrontal cortex leads to improved statistical learning, it would raise the possibility of a right-hemisphere gating mechanism, in line with the classic HERA model.
2. Our study was designed to test competing hypotheses about the DLPFC’s gatekeeping function – its role in controlling access to the long-term memory systems that store prior knowledge and internal models. Based on this hypothesis, we predict enhanced statistical learning will be accompanied by impaired episodic retrieval in the stimulation groups. If, however, DLPFC inhibition affects statistical learning independently of episodic memory, it would suggest that the DLPFC’s influence on statistical learning is mechanistically distinct from its role in accessing the model-based priors required for episodic retrieval.

## Results

### Is there a difference between the groups in statistical learning? Reaction time analysis

We used the Alternating Serial Reaction Time (ASRT) task to assess statistical learning. In this task, learning is signified by the performance difference between high- and low-probability triplets: if participants react faster to stimuli that belong to a triplet that occurs with a high probability as compared to triplets that occur with a low probability, it means they have an implicit knowledge of the underlying regularity. Importantly, the relevant effect is not the absolute RT (and later, accuracy) difference between groups, but the between-group difference in the high-versus low-probability triplet contrast.

We performed linear mixed-effect models both using frequentist and Bayesian methodology (see Methods for details). The final model in both cases included only the random intercept by participant (RT ∼ Group × Epoch × Triplet type + (1| Participant)). For the former one, model details are shown in Supplementary Table S1. For the latter one, we can draw conclusions based on the credible interval (if it includes zero, it means we do not have a strong evidence for the effect), we report the relevant comparisons in the Supplementary Materials, see the exact table number below in the text. See Figure 1 for details of the RT results.

**Figure 1.**
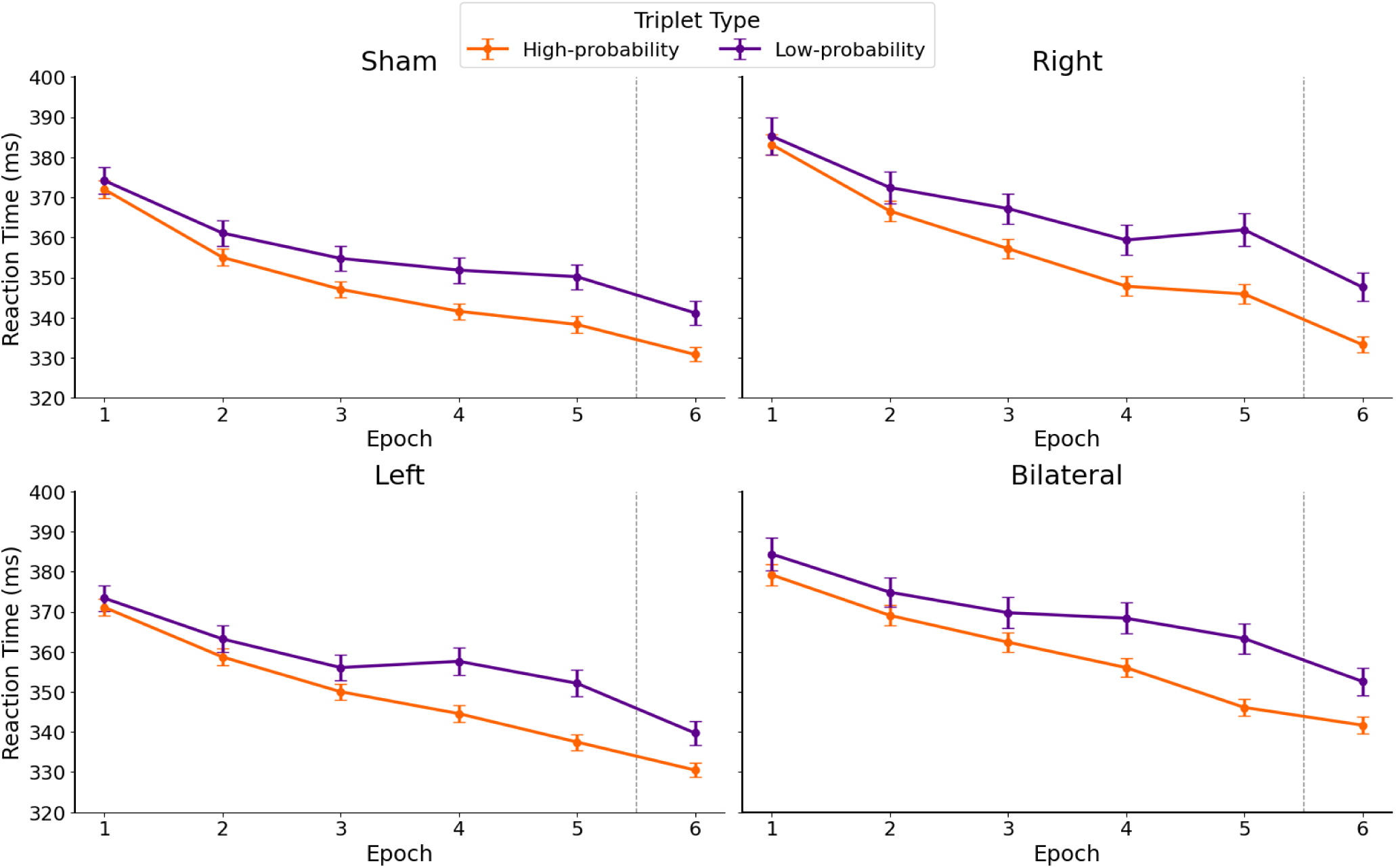
Reaction times to high- and low-probability triplets in sham, right, left, and bilateral stimulation groups. The figure displays reaction times (in milliseconds) for high-probability (orange) and low-probability (purple) triplets across six epochs (x-axis). The panels (arranged from top right to bottom left) correspond to the sham, right, left, and bilateral DLPFC stimulation groups. The difference in reaction times between high- and low-probability triplets indicates statistical learning. The dashed grey line between the 5th and 6th epochs marks the 24-hour interval between the two experimental sessions. Error bars represent 95% confidence intervals of the mean reaction time.

The frequentist linear mixed-effect model on RT revealed a significant main effect of Epoch, indicating that participants became faster throughout the task, regardless of triplet types (*F*(5, 183988.17) = 728.44, *p* < 0.001). This general improvement differed between the groups, as reflected by a significant Group × Epoch interaction (*F*(15, 183988.17) = 4.02, *p* < 0.001). The post hoc comparisons showed that the sham and the right group improved in RT in all epochs, except between the 4th and 5th epochs (p < 0.027 in all consecutive epoch pairs besides 4-5th, where p = 0.539), where they stagnated. The left group showed a similar pattern, except that they did not improve between the 3rd and 4th epochs (p = 0.795), but did so between all others (p < 0.001 in all other consecutive epoch pairs). The bilateral group improved during the whole task (p < 0.044). A significant main effect of Triplet Type was also observed (*F*(1, 183988.40) = 486.98, *p* < 0.001), with faster responses to high-probability triplets than to low-probability ones indicating statistical learning. The difference between the Triplet Types changed over time, shown by the Epoch × Triplet Type interaction (*F*(5, 183988.32) = 17.54, *p* < 0.001), signifying learning improved over time. There was no significant Group main effect, indicating that the groups showed similar overall RT if we disregard triplet types (*F*(3, 91.13) = 0.88, *p* = 0.453).

Most importantly, we found a significant Group × Triplet Type interaction (*F*(3, 183988.40) = 3.71, *p* = 0.011), showing that the magnitude of the statistical learning effect (i.e., the RT difference between high- and low-probability triplets) varied across groups. The right stimulation group demonstrated significantly greater statistical learning than the sham group (*p* = 0.005, SE = 1.14) and the left stimulation group (*p* = 0.032, SE = 1.17). Additionally, the bilateral stimulation group outperformed the sham group (*p* = 0.015, SE = 1.12). A trend-level difference was also observed between the bilateral and left stimulation groups, with the bilateral group showing better learning (*p* = 0.079, SE = 1.15). Full post hoc pairwise comparisons, including effect estimates and 95% confidence intervals, are reported in Supplementary Table S3. This Group × Triplet Type interaction is illustrated in Figure 2. However, this difference between the groups in statistical learning did not change over time, as the Epoch × Triplet Type × Group interaction was not significant (*F*(15, 183988.32) = 0.53, *p* = 0.927). The model’s explanatory power was modest (Marginal R² = 0.029, Conditional R² = 0.171).

**Figure 2.**
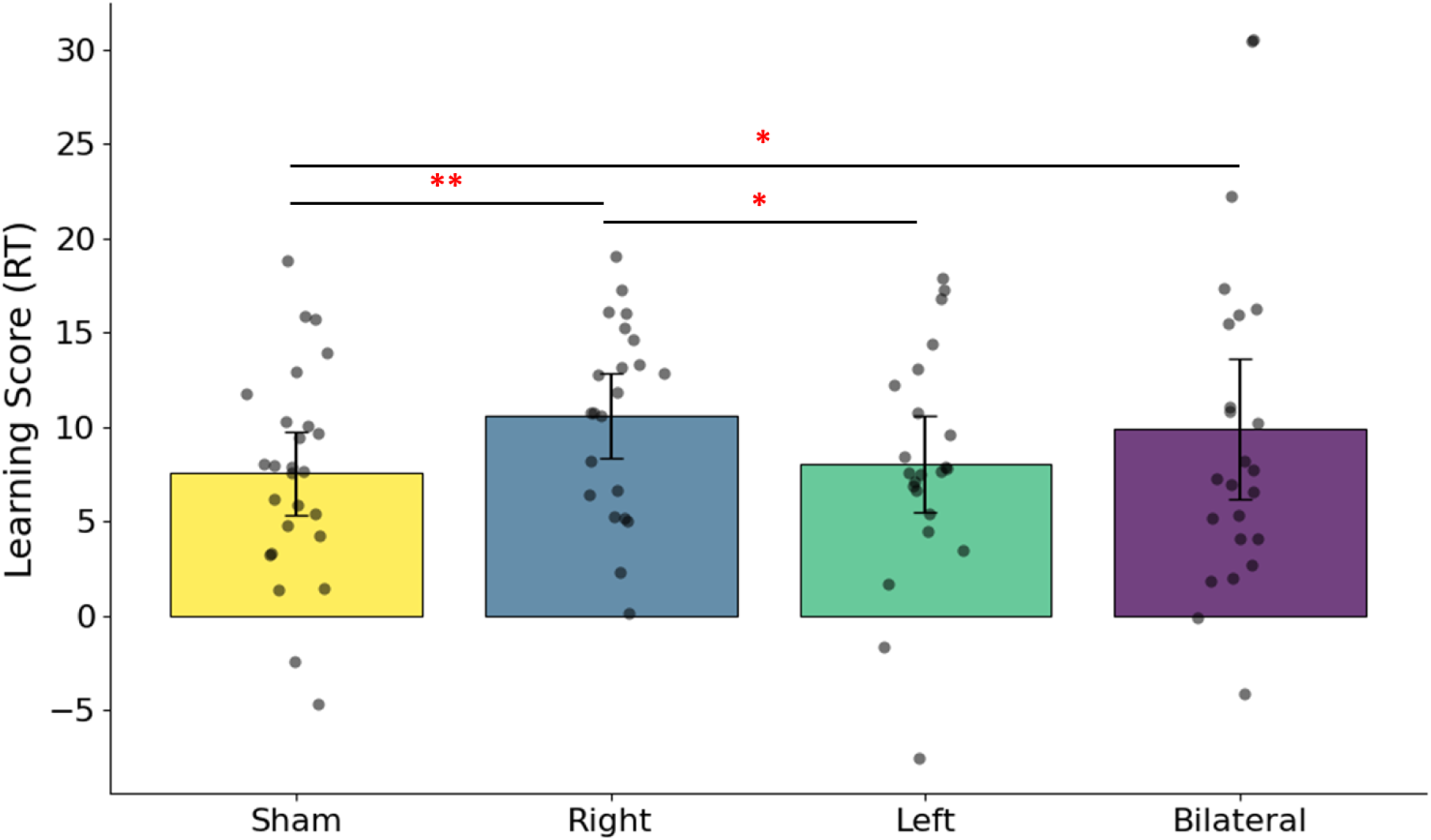
Group differences in reaction time (RT) learning score. Learning scores were computed as the difference between RT to low- and high-probability triplets (RT low − RT high) and averaged within each stimulation group (sham, right, left, bilateral). Higher values reflect stronger statistical learning. Bars represent group mean learning scores, and error bars indicate 95% confidence intervals of the mean. Asterisks indicate statistical significance (*p < 0.05, \*\**p < 0.01)*.

The Bayesian model provided evidence for the Epoch main effect (see Supplementary Table S6), supporting the frequentist results. It failed to be decisive regarding the Group main effect and the Group × Epoch interaction (Supplementary Table S8 and S7, respectively), thus, we cannot draw strong conclusions regarding potential group differences on statistical learning-independent RT. There was evidence for the Triplet Type main effect (Supplementary Table S5), and the Epoch × Triplet Type interaction (Supplementary Table S9), which supports the existence of statistical learning and its improvement throughout the task. Most importantly, the Bayesian evidence supported our main result: there was a reliable difference between the right and the sham, the bilateral and the sham, and the right and the left groups, see Supplementary Table S10. The three-way interaction was uncertain according to the Bayesian model (Supplementary Table S11).

### Is there a difference between the groups in statistical learning? Accuracy analysis

Similarly to our RT analyses, to assess learning differences as measured by accuracy, we performed both frequentist and Bayesian generalized linear mixed-effect models (see Methods for details). For the former, model details are shown in Supplementary Tables S2 and S4. For the latter, model convergence was satisfactory for all parameters (R̂ ≤ 1), with effective sample sizes being adequate for both bulk and tail posterior distributions. The credible intervals of the Bayesian analyses are shown in Supplementary Table S12. In both cases, the best model included the random intercept per participant (Accuracy ∼ Group × Epoch × Triplet type + (1| Participant)). Figure 3 illustrates the time course of accuracy across the stimulation groups.

**Figure 3.**
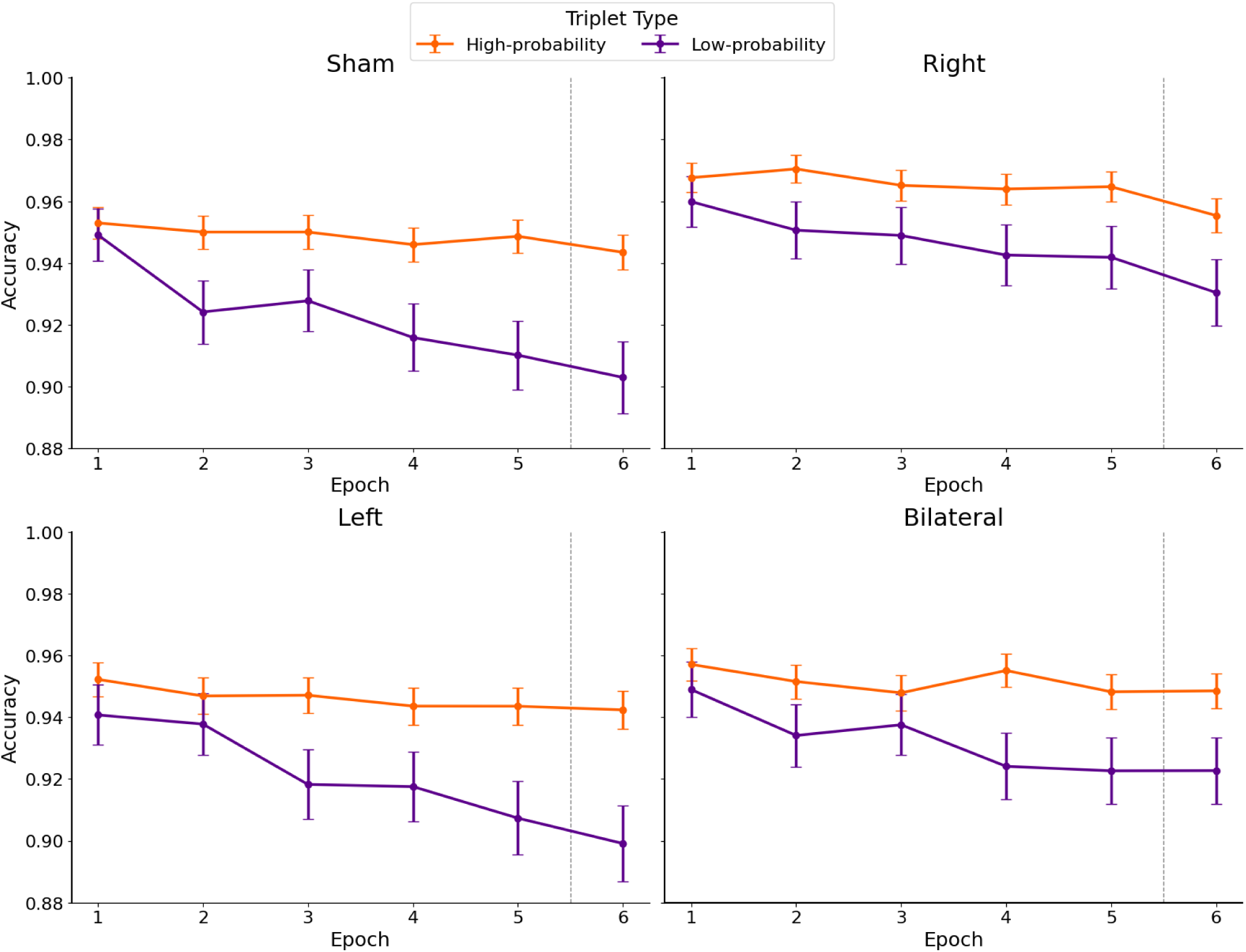
Accuracy to high- and low-probability triplets in sham, right, left and bilateral stimulation groups. The figure displays accuracy for high-probability (orange) and low-probability (purple) triplets across six epochs (x-axis). The panels, arranged clockwise from the top left, correspond to the groups as follows: top-left – sham; top-right – right stimulation; bottom-right – bilateral stimulation; bottom-left – left stimulation. The difference between high- and low-probability triplets serves as an index of statistical learning. The dashed grey line between the 5th and 6th epochs marks the 24-hour interval between the two experimental sessions. Error bars represent 95% confidence intervals of the mean accuracy.

The frequentist generalized linear mixed model (GLMM) on accuracy (ACC) revealed a significant main effect of Epoch, *χ²*(5) = 141.50, *p* < 0.001, indicating that participants became more accurate at the beginning of the task (Epoch 1–2) and less accurate in the later phases (Epoch 4–5), consistent with practice and fatigue effects. A robust main effect of Triplet Type was also observed, *χ²*(1) = 363.95, *p* < 0.001, with higher accuracy for high-probability compared to low-probability triplets, indicating statistical learning. The main effect of Group was not significant, *χ²*(3) = 5.46, *p* = 0.141, suggesting no overall group differences in accuracy. Similarly, there were no significant two-way interactions between Group and Epoch, *χ²*(15) = 13.69, *p* = 0.549. There was no significant interaction between Group and Triplet Type, *χ²*(3) = 2.53, *p* = 0.470. In contrast, the Epoch × Triplet Type interaction was significant, *χ²*(5) = 32.35, *p* < 0.001, suggesting that the magnitude of statistical learning fluctuated across different time points. However, the three-way Group × Epoch × Triplet Type interaction was not significant, *χ²*(15) = 14.48, *p* = 0.489, indicating no reliable group-specific dynamics in the evolution of statistical learning.

The Bayesian model supported the Epoch main effect, showing a decrease in accuracy between Epoch 1 and 2, see Supplementary Table S13. Moreover, in contrast with the frequentist results, the model provided evidence for the Group main effect and the Group × Epoch interaction, suggesting that the right group has performed more accurately (regardless of triplet types) compared to the left group (Supplementary Table S14), which was driven by a group difference between the right and the left group in Epoch 2-5 (Supplementary Table S15). This and the uncertain results on RT raise the possibility of group differences in speed-accuracy trade-off. To rule this out as an explanation to our main results (concerning statistical learning), we performed additional analyses reported in the Supplementary Results.

The model supported the Triplet type main effect and the Epoch × Triplet Type interaction, providing evidence for statistical learning and its development across epochs (See Supplementary Tables S12 and S17, respectively). There was no reliable evidence on the Group × Triplet Type interaction (Supplementary Table S16). Bayesian evidence was largely not provided for the Group × Epoch × Triplet Type interaction either, except for sporadic results of a sham-left difference in Epoch 1-Epoch 2 contrast and left-bilateral difference in the Epoch 2-Epoch 3 contrast, see Supplementary Table S18.

The complete outputs of the RT and ACC models are provided in Supplementary Table S1 and S2. Model-estimated reaction times and accuracy for high- and low-probability triplets across stimulation groups and epochs (for the frequentist model) are presented in Supplementary Figures S1 and S2.

### Do the groups differ in episodic retrieval?

In item memory, there was a significant main effect of epoch (*F*(2.27, 197.29) = 30.01, *p* < 0.001, *η²p* = 0.256, *BF_incl_* > 2.47×10^17^), indicating that item memory scores decreased across time (forgetting across epochs, *p* `≤ 0.015). The Epoch × Group interaction was not significant` (*F*(6.80, 197.29) = 0.66, *p* = 0.70, *η²p* = 0.022, *BF_incl_* = 0.073), suggesting that changes over time did not differ significantly between groups. The main effect of Group was also not significant (*F*(3, 87) = 1.12, *p* = 0.344, η²p = 0.037, *BF_incl_*= 0.018), indicating a lack of significant group differences in item memory. The results are presented in Figure 4 panel A. The automatic association scores showed a similar pattern: the significant Epoch main effect (*F*(2.08, 181.11) = 4.07, *p* = 0.017, *η²p* = 0.045, *BF_incl_* = 3.595), indicated forgetting in the second session (i.e., the epoch after the 24h delay) compared to the last epoch of the first session, however, this was reduced to a trend after correcting for multiple comparisons (*p* = 0.092). Neither the Epoch × Group interaction (*F*(6.24, 181.11) = 0.80, *p* = 0.574, η²p = 0.027, *BF_incl_*= 0.045), nor the Group main effect (*F*(3, 87) = 1.07, *p* = 0.367, *η²p* = 0.035, *BF_incl_* = 0.032), reached the level of significance, indicating no group differences in the overall score, or in the course of forgetting. See Figure 4 panel B for details. In the recollection score analysis, the main effect of Epoch was not significant (*F*(3, 261) = 1.94, *p* = 0.124, *η²p* = 0.022, *BF_incl_* = 0.198), indicating no reliable change in recollection scores over time. The Epoch × Group interaction was not significant (*F*(9, 261) *=* 0.72, *p* = 0.690, *η²p* = 0.024, *BF_incl_* = 0.034), suggesting that any temporal changes did not differ between groups. The main effect of Group, similarly to the other scores, was not significant (*F*(3, 87) = 0.71, *p* = 0.549, *η²p* = 0.024, *BF_incl_* = 0.024). The recollection results are shown on Figure 4 panel C.

**Figure 4.**
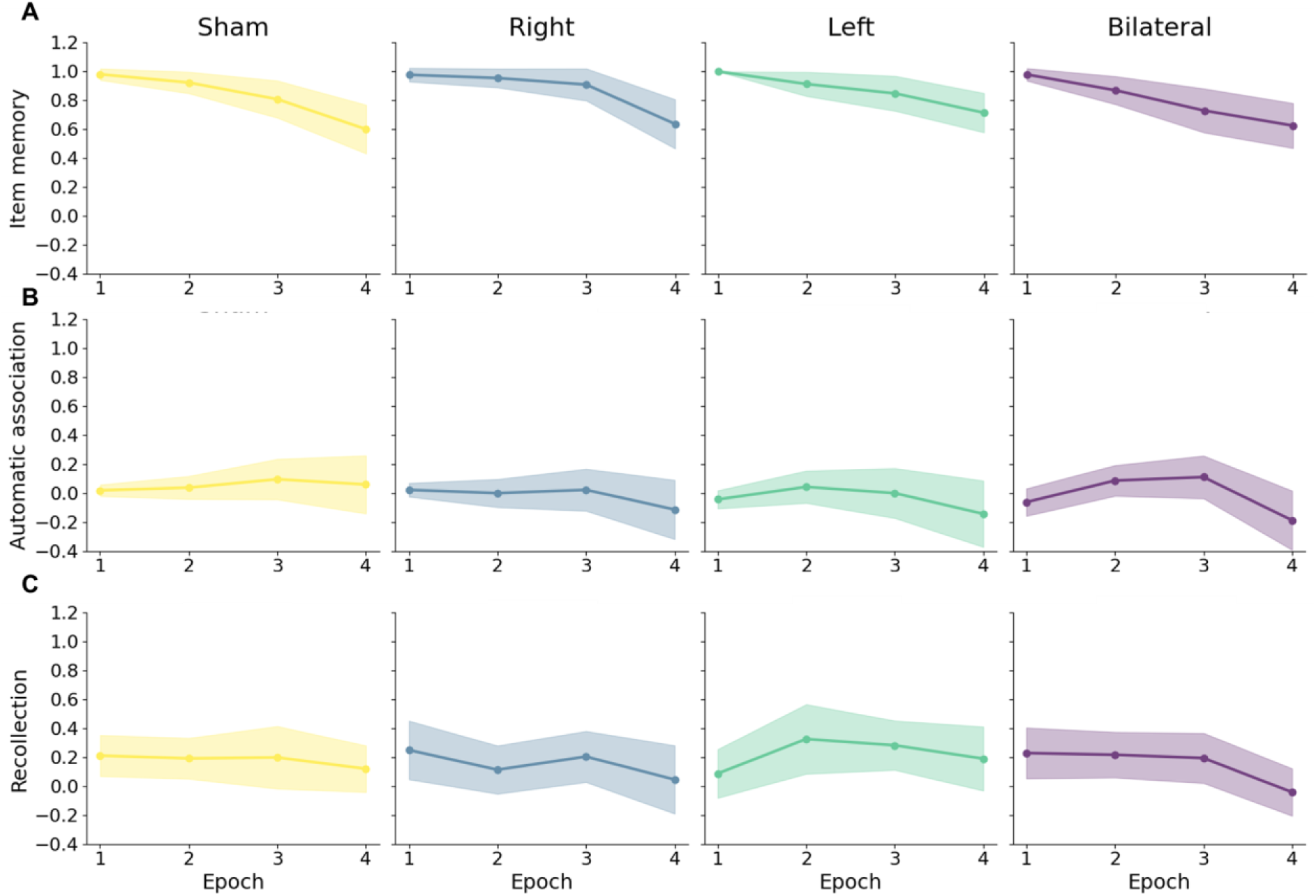
Paired Associate Learning Task results. The y-axis reflects the three different PALT scores, x-axis show the epoch when the data was collected (please note that in Epoch 2 and 4, participants did not perform the PALT task). Colors indicate the stimulation groups. Error bands show the 95% confidence intervals. Panel A): Item memory score on the y-axis. Panel B): Automatic association score on the y-axis. Panel C): Recollection score on the y-axis.

### Exploratory analyses: individual differences

#### Does the variance of reaction times differ due to the stimulation?

To test whether participants in the stimulation groups showed a different variability of reaction times, we performed a Levene’s test with group (sham/right/left/bilateral DLPFC stimulation) as the independent variable. Levene’s test indicated significant differences in variance across groups [*F*(3, 184123) = 270.65, *p* < 0.001]. Results were confirmed using a Brown–Forsythe test, yielding the same pattern of group differences. The right and the bilateral DLPFC stimulated groups showed significantly higher variances compared to the sham and the left DLPFC stimulated groups (*p_Holm_* < 0.001 in each case). For easier interpretability, we report the standard deviations (rather than the variance) in Table 1 and Figure 5.

**Figure 5.**
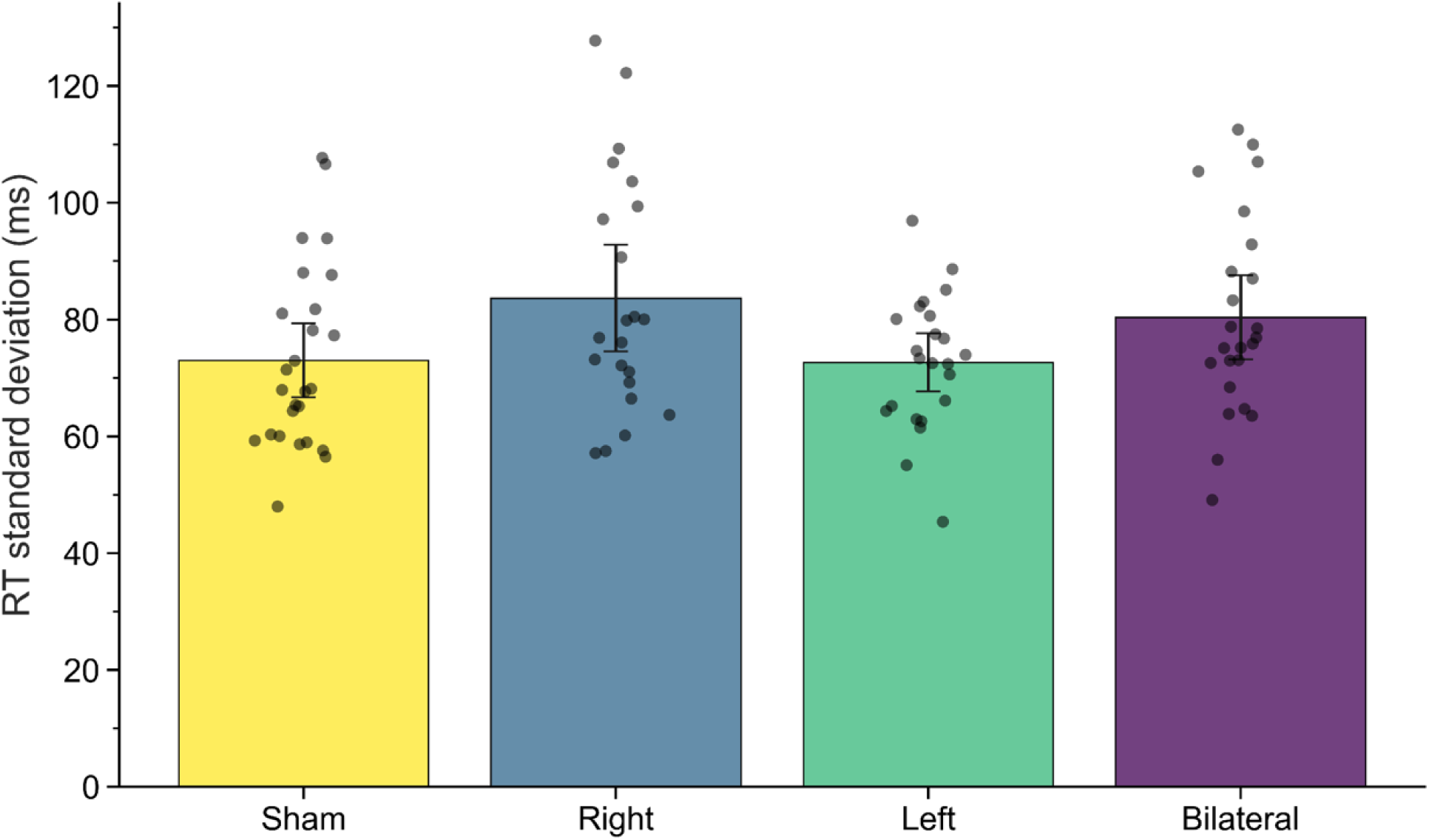
The RT standard deviations (y-axis). The x-axis and the colors depict the four groups. Black dots represent individual RT standard deviation data.

**Table 1.**
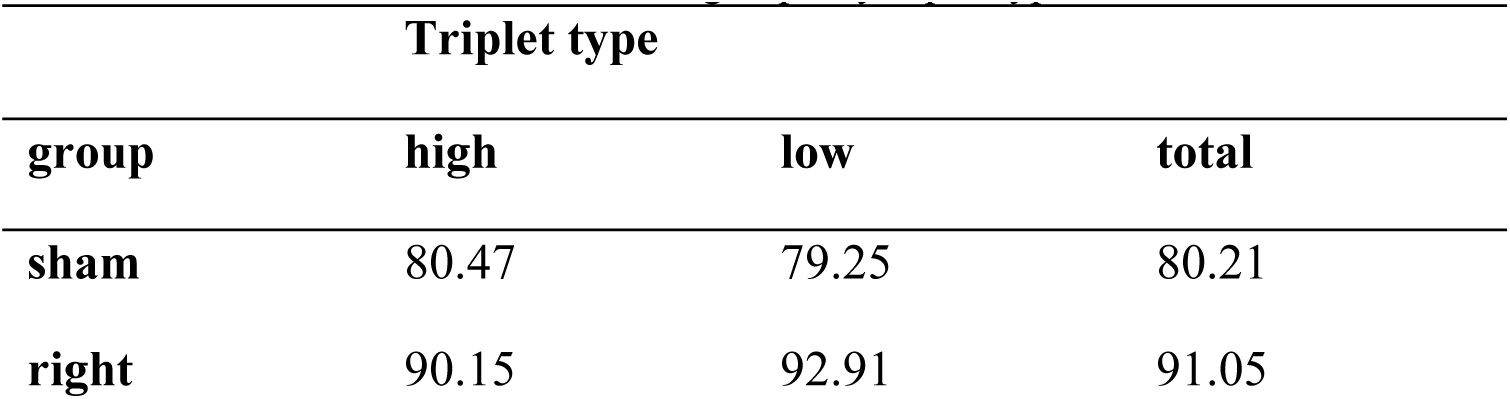
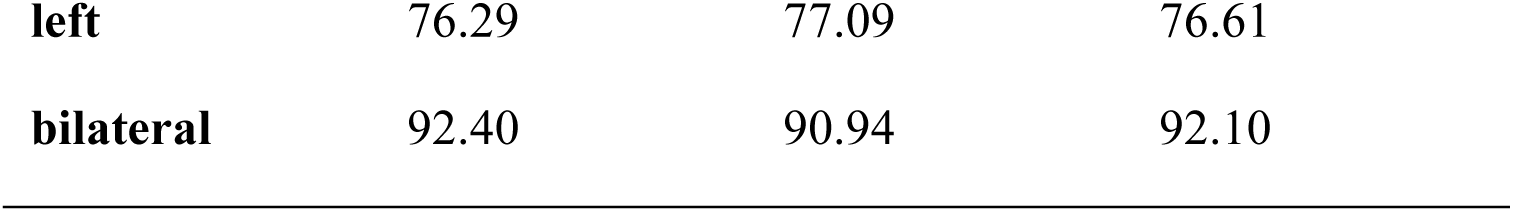
RT standard deviations of the groups, by triplet type.

| Triplet type |  |  |  |
| --- | --- | --- | --- |
| group | high | low | total |
| sham | 80.47 | 79.25 | 80.21 |
| right | 90.15 | 92.91 | 91.05 |
| **left** | 76.29 | 77.09 | 76.61 |
| **bilateral** | 92.40 | 90.94 | 92.10 |

#### Does working memory correlate differently with statistical learning across groups?

We tested the correlation between statistical learning (defined as the RT difference of low- and high-probability triplets throughout the whole task) and CSPAN, separately for each group. We found no significant correlation in any of the groups, although the bilateral DLPFC stimulated group showed a trend-level positive correlation [sham: *r*(24) = -0.068, *p* = 0.741; right: *r*(20) = -0.223, *p* = 0.296; left: *r*(21) = -0.089, *p* = 0.686; bilateral: *r*(22) = 0.369, *p* = 0.076], see Figure 6.

**Figure 6.**
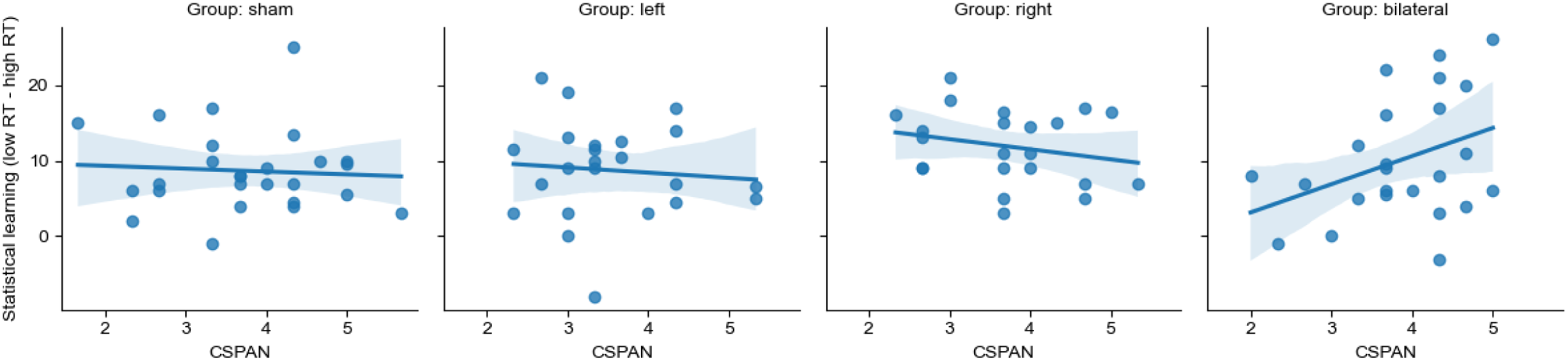
Statistical learning as a function of baseline working memory capacity. The y-axis represents the RT difference of low-minus high-probability triplets (calculated for the sake of this analysis alone). The x-axis represents the Counting span (CSPAN) performance. Dots indicate individual data, lines represent a linear regression trend line, and bands show the 95% confidence interval of the regression estimate.

## Discussion

We aimed to causally assess the role of the dorsolateral prefrontal cortex (DLPFC) in statistical learning – an unsupervised form of learning critical for extracting regularities from sensory input, acquiring new skills, and constructing predictive representations – by applying inhibitory transcranial magnetic stimulation (rTMS) over the left, right, or bilateral DLPFC. We found that inhibition of the right and bilateral DLPFC significantly enhanced statistical learning performance, as indicated by faster responses to predictable compared to unpredictable stimuli. To examine the functional specificity of this effect, we included an episodic memory task as a cognitive control measure. Episodic memory retrieval remained unaffected across all stimulation groups, suggesting that the observed enhancement in statistical learning cannot be attributed to a general facilitation or disruption of cognitive performance. Together, these findings demonstrate a right-lateralized influence of the DLPFC on statistical learning, suggesting that this effect may operate through mechanisms distinct from episodic retrieval as measured here. Crucially, the selective enhancement of statistical learning in the right and bilateral groups was observed in reaction times, while statistical learning measured by accuracy remained stable across all groups. This lack of a Group × Triplet Type interaction in accuracy suggests that the observed effects reflect a specific modulation of the processing of statistical regularities rather than a generalized performance decrement, increased response noise, or a simple speed-accuracy trade-off.

Our findings in the bilateral stimulation group are in line with the only prior rTMS study on the acquisition phase of statistical learning using bilateral DLPFC inhibition, which similarly reported enhanced learning of probabilistic regularities. One additional study has applied bilateral inhibitory stimulation during the retrieval phase and found improved retrieval of statistical information^16^. These findings provide converging causal evidence for the involvement of the DLPFC in statistical learning, contributing to a growing literature suggesting an inverse relationship between DLPFC-mediated functions –-such as executive control –- and statistical learning performance^7,8,15,29,30^. This inverse relationship supports the competition theory, which posits that cognitive control processes –-although beneficial for goal-directed, model-based processes –-interfere with the automatic, model-free mechanisms underpinning statistical learning. Evidence from functional connectivity and developmental studies further supports this interpretation: reduced functional coupling between the DLPFC and other brain regions has been associated with improved statistical learning^31,32^; and the learning advantage observed in children may stem from the immaturity of the DLPFC^10–12,33^. Collectively, these findings suggest that reduced prefrontal engagement, whether through development or neuromodulation, can facilitate statistical learning by releasing the system from top-down constraints.

We sought to investigate whether the apparent competition between DLPFC functioning and statistical learning could be explained by the DLPFC’s hypothesised control over the access to long-term memory (i.e., its gating role)^9^. Working memory plays an important role in accessing long-term memory and is often conceptualized as a state of activated long-term memory^26^. Since our 1 Hz rTMS stimulation targeted a major working memory hub^34,35^, it is reasonable to assume that we also influenced access to long-term memory. Based on prior theoretical proposals^9^, we hypothesized that disrupting DLPFC activity may impair access to pre-existing internal models stored in long-term memory, thereby facilitating the acquisition of novel statistical regularities. While our findings on statistical learning support this hypothesis, we did not reveal a disruption of episodic retrieval following DLPFC inhibition. Thus, we did not find evidence that this effect was mediated by episodic memory interference. Alternative gate pathways remain plausible, or gating may happen through different memory systems, like semantic or working memory. Overall, while our results do not provide support for a gating mechanism based on episodic memory interference, they leave open the possibility that the DLPFC plays a modulatory role through alternative routes - a hypothesis that warrants further investigation.

Our hemispheric findings offer a more specific interpretation. A plausible explanation for our results lies in the modality-specific lateralization of prefrontal cortex function. Our use of a visuomotor statistical learning task aligns perfectly with findings that right DLPFC disruption primarily affects visual learning^21,22^. This framework also helps reconcile seemingly conflicting results in the literature, which have often reported enhanced *verbal* statistical learning following *left* DLPFC disruption^15,29^. This right-lateralized effect is also broadly consistent with the Hemispheric Encoding/Retrieval Asymmetry (HERA) model^27,28^. This model posits a primary role for the right DLPFC in memory retrieval. From this perspective, inhibiting the right DLPFC may have impaired access to pre-existing models, shifting the cognitive system toward more data-driven processing and thereby facilitating statistical learning. The results on the right hemisphere-specific effect are consistent with the idea that our episodic memory task lacked the sensitivity to detect subtle group differences (see limitations), and that statistical learning was nonetheless improved via an episodic gating mechanism^9^ –-future studies are needed to test this speculation. Taken together, our results provide causal evidence that right DLPFC inhibition enhances visuomotor statistical learning, likely by attenuating top-down, memory-based constraints. Future studies using systematic, modality-sensitive designs are needed to further elaborate this relationship.

To further interpret our findings, it is essential to consider which cognitive strategies and information processing styles are known to support statistical learning. While frequency-maximizing and frequency-matching strategies were discussed in the introduction^4,19^, they are unlikely to explain our results, as such strategies would predict enhanced learning following left DLPFC inhibition, which we did not observe. An alternative potential mechanism draws from reinforcement learning research: trying new, undiscovered patterns (that is, exploration as opposed to exploitation) and a broader, more distributed attention to various environmental signals (that is, broader information sampling) can facilitate the early acquisition of probabilistic regularities^36,37^, as frequent re-sampling could lead to a deeper knowledge of statistical structures that are inherently probabilistic (for a discussion, see ref^38^). Although our study was not specifically designed to assess differences in exploration/exploitation or their causal relationship with statistical learning, we propose that right and bilateral DLPFC inhibition fostered a shift toward a more exploratory information-sampling strategy. This interpretation is supported by our finding of significantly greater RT variability –-a well-established behavioural proxy for exploration^39–41^ –-in the same groups that showed enhanced learning. We speculate that the observed benefits after right and bilateral DLPFC inhibition in statistical learning may reflect a shift toward a broader, more exploratory information processing style.

The observed shift toward broader, more exploratory information sampling may be caused by the disruption of working memory: lower DLPFC functioning and thus lower working memory have been linked to such a strategy^42,43^. Individuals with reduced working memory capacity (such as children or participants undergoing DLPFC inhibition in our study) tend to re-sample their environment more frequently, as previously acquired information is more susceptible to decay^42^. This explanation is compelling because our stimulation target, the Brodmann 9 area, is known for its role in working memory^35,44^, and it is consistent with extensive evidence that working memory exerts a modulatory influence on statistical learning^7,8,21^. This interpretation is further strengthened by a recent work indicating that TMS over the DLPFC facilitates inhibitory firings in executive control-related networks on the cell-level^45^. This framework may also reconcile the above-mentioned conflicting findings regarding the lateralization of statistical learning. Inhibiting verbal working memory, which shows a well-established left-lateralization^46–49^, may result in a broader information sampling strategy in the verbal domain^50^, benefiting verbal statistical learning tasks^15,29^. But in our visuomotor statistical learning task, verbal working memory (and thus, broader verbal information sampling) could have little effect, hence the null result in the left stimulation group. On the other hand, right stimulation may have been effective because it influenced visual working memory (known to be right-lateralized^44,46–49^), thus shifting visual processing to be more explorative. While our study was not designed to directly test this hypothesis, exploratory analyses offer tentative support. Specifically, we observed a trend-level positive correlation between working memory capacity and statistical learning in the bilateral group, suggesting that DLPFC inhibition may enhance statistical learning, particularly in individuals with higher baseline working memory capacity –-showing the baseline working memory’s relation to the stimulation effectiveness (see Figure 5). Taken together, we speculate that DLPFC inhibition may reduce working memory maintenance of prior samples, promoting increased environmental exploration, which facilitates statistical learning of probabilistic regularities –-future studies should be designed specifically to probe this mechanism.

Collectively, our findings fit within recent theoretical frameworks that challenge area-centric models of cognition in favor of distributed, brain-wide dynamics. Hayden and colleagues^51^ argue that the "arealization paradigm" – treating cortical regions as stable, modular units – obscures the context-dependent nature of neural computation. In parallel, Rosen and Freedman^52^ suggest that while cognitive signals are broadly reflected across the brain, they are not ubiquitous, appearing instead in task-dependent, reconfiguring networks. Our results operationalize these ideas: inhibitory rTMS over the DLPFC did not eliminate a categorical function but instead dynamically reweighted the balance of processing within a large-scale network. The selective enhancement of statistical learning following right and bilateral inhibition – coupled with increased reaction time variability and preserved episodic memory – is inconsistent with strict localization or simple gating accounts. Rather, reducing right-prefrontal top-down constraints likely shifted the system toward a more exploratory, distributed, model-free state through affecting larger networks, rather than solely the DLPFC. This demonstrates that cognitive outcomes reflect the dynamic reweighting of large-scale networks rather than the isolated engagement of discrete areas, providing a causal bridge between neuroanatomy and functional flexibility.

A key limitation of our study is our choice of task for episodic memory retrieval. We found reliable null effects on episodic retrieval regarding the stimulation effects, which were supported by Bayesian evidence. Although it is not without example to find no DLPFC-related effect on episodic memory, the majority of studies were able to show a relation to the DLPFC^25^. This may have multiple explanations. As we could see earlier, DLPFC lateralization greatly depends on the modality of the task in many cognitive functions, and episodic memory is no exception^27,28,53^. Our task had both visual and verbal elements, enabling participants to use their preferred strategy, either verbal or visual. Thus, this task was a suboptimal choice for a study that wanted to explore lateral effects. However, compensation from the non-inhibited strategy (verbal in the case of left inhibition, visual in the case of right inhibition) does not account for our results in the bilateral group: when both hemispheres are inhibited, we should have seen the effect if it were there. Furthermore, the PALT measures a specific form of associative memory. It is therefore possible that the DLPFC’s gating function does not act through this declarative retrieval but rather through other forms of memory that are more integral to the online processing of statistical information. For instance, DLPFC inhibition may have disrupted access to semantic representations or the short-term maintenance of statistical priors within working memory itself –-cognitive processes not captured by our episodic task. Nevertheless, the question arises whether our task, and the power in our study was sufficient to capture group differences. The task was able to capture memory changes per se, demonstrated by the item memory and automatic association scores reliably showing expected temporal dynamics (i.e., forgetting across epochs in item memory).

We acknowledge that recollection failed to sensitively reflect the expected dynamics, thus, this score should be interpreted with caution. We would like to outline a possible explanation here. Recollection (as per our definition) can be supported by two distinct neural patterns: temporal activation (among others, in the hippocampus and medial PFC) when the recollection happens, and sustained activity (among others, in the striatum and left inferior frontal gyrus) that lasts throughout the delay^82,83^. In our design, participants were distracted throughout the delay by the ASRT task, thus, this sustained activity was unlikely to last until the retrieval phase. As item memory is often utilized when recollection fails^84^, it is possible that our task triggered item memory to a greater extent, while participants did not rely strongly on the controlled recollection of item pairs. If this explanation stands, it means that our null results on recollection reveal more on the nature of episodic memory, rather than on the effect of the stimulation, thus, have to be interpreted with caution. This, however, does not question the results on item memory and automatic association. Moreover, the Bayesian analyses on all three metrics provided moderate-to-strong evidence against group differences, thus, the null results are unlikely to derive from power issues. Future studies should systematically explore the effect of modality and test various memory systems in relation to statistical learning to clarify the precise nature of the DLPFC’s modulatory role.

The present study employed a stimulation methodology consistent with prior work (e.g. ref^9,16,54–62^). Nevertheless, several methodological refinements could further increase anatomical precision and control for non-specific stimulation effects in future investigations. For example, incorporating MRI-guided neuronavigation would reduce inter-individual anatomical variability and allow more precise characterization of the stimulated subregion within PFC^63^. Similarly, employing an active control site within participants, a dedicated sham coil, or alternative sham protocols that better reproduce somatosensory aspects of stimulation would provide a stronger control for non-specific effects. While we acknowledge that these methodological improvements could have further strengthened our results, they do not fundamentally question the interpretation of our results. Several aspects of the present findings argue against a primarily non-specific or placebo-based explanation. First, the two lateral stimulation groups essentially served as active controls for one another: if the effects were driven mainly by generic stimulation-related factors, similar patterns would be expected across hemispheres. Instead, we observed significantly differing effects in the left and right stimulation groups, which supports functional specificity. Second, although EEG-cap–based targeting does not offer the spatial precision of neuronavigation, it is widely used and has been shown to produce reliable behavioural modulation ^64^. Importantly, any residual spatial variability would be expected to reduce sensitivity rather than systematically produce condition-specific or opposing effects^65^. Finally, the same stimulation protocol that modulated statistical learning did not influence episodic memory performance, and Bayesian analyses provided evidence in favor of the null effect in that domain. Together, these considerations suggest that while future studies using complementary targeting and sham approaches would further strengthen confidence in the anatomical specificity of the effect, the present results are unlikely to be attributable to non-specific stimulation or targeting imprecision.

In summary, our study provides novel causal evidence for a right-lateralized role of the DLPFC in modulating statistical learning. These findings offer support for the competition theory by demonstrating that DLPFC-mediated functions, particularly those localized to the right hemisphere, can suppress model-free learning processes. Notably, our results do not support the interpretation that this effect arises from reduced interference by episodic memory systems. Instead, they suggest that, if such a gating mechanism exists, it operates independently of episodic retrieval. It raises the possibility of a long-term memory gating mechanism specifically for unsupervised statistical information. Furthermore, we propose an alternative account in which DLPFC inhibition alters information sampling strategies, promoting a broader, more exploratory mode of learning. The observed increase in RT variability, alongside the hemispheric specificity of the effects, supports this view and highlights a potential modality-dependent component in the prefrontal contribution to statistical learning. Future research should more directly investigate the mechanisms linking working memory capacity, information sampling style, and statistical learning. Our study provides insight and proposes novel hypotheses that will be critical for advancing our understanding of the DLPFC’s interaction with statistical learning across both typical and altered brain states.

## Materials and Methods

### Participants

We recruited 104 healthy adults to our experiment with normal or corrected-to-normal vision. They were recruited through a university course and received course credits for their participation. Inclusion criteria were no history of neurological or psychiatric diagnoses and no contraindication for non-invasive brain stimulation (epilepsy in the participant or their first-degree relatives, metal implants in the head or neck, pacemaker, migraine, history of severe head injury or concussion). All of them provided written informed consent. We excluded eight participants due to technical issues, and one participant who did not return for the second session. Additionally, 4 participants were excluded from the PALT analysis as they did not finish the task on the second day due to technical issues. Thus, in the ASRT analyses, we entered the data of 95 participants. The experiment was conducted in accordance with the Declaration of Helsinki, with ethical approval granted by the Ethics Committee of Eötvös Loránd University (approval number: 2019/282).

As detailed in the Procedure section below, participants were randomly assigned to one of four experimental groups: (1) active right DLPFC stimulation, (2) active left DLPFC stimulation, (3) active bilateral DLPFC stimulation, or (4) sham stimulation (control group). Demographic information and complex working memory (counting span, CSPAN) of the participants are shown in Table 2. There were no significant differences between the groups in terms of gender, age, or CSPAN performance.

**Table 2.**
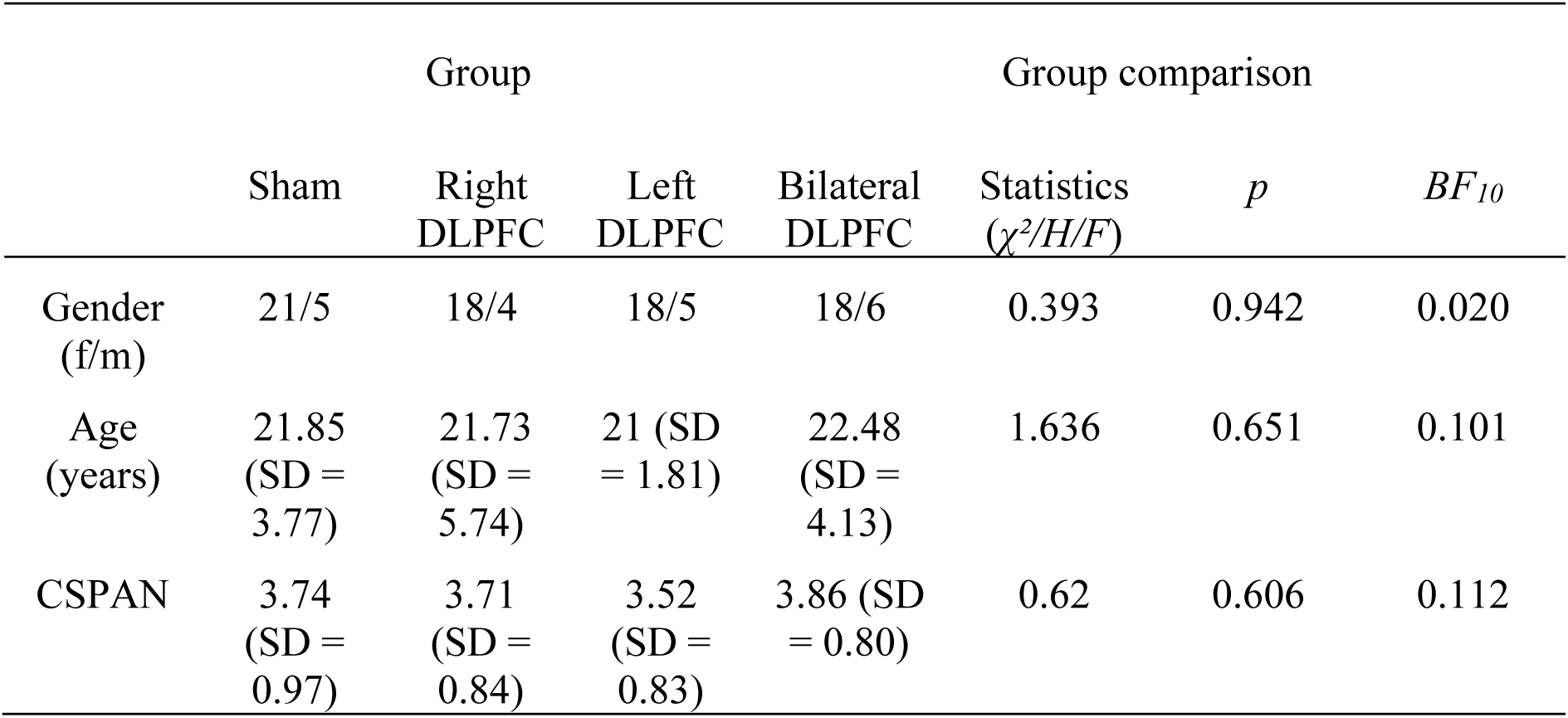
Demographic characteristics and counting span (CSPAN) of the experimental groups.

### Tasks

#### Alternating Serial Reaction Time Task

To measure statistical learning, we employed the Alternating Serial Reaction Time (ASRT) task^66^. In this task, four black circles were displayed horizontally on a white background of the computer screen, with dog head stimuli appearing one at a time inside these circles (see Figure 6, panel A). Each circle corresponded to a specific response key (Y, C, B, M) on a Hungarian QWERTZ keyboard layout. Participants were instructed to press the appropriate key as quickly and accurately as possible after the stimulus appeared, using their index and middle fingers of both hands. The stimulus remained visible until the correct key was pressed, after which the next stimulus appeared following a 120 ms response-to-stimulus interval.

The task was organized into ∼1-minute-long blocks. Each block began with five random stimuli (for practice purposes), which were excluded from later analyses. Following these, an eight-element sequence was repeated ten times, resulting in a total of 85 trials per block. Within this repeating sequence, every first element was a random stimulus, while the alternating elements followed a fixed, predetermined sequence (e.g., 2–r–4–r–3–r–1–r, where numbers denoted the fixed pattern elements, and "r" represented a randomly selected position from the four locations). Importantly, participants were unaware of the embedded sequential regularity. The alternating pattern of fixed and random stimuli generated triplets - runs of three elements formed by a fixed structure of stimulus transitions - in a sliding window manner. Based on the example above, triplets such as 2_X_4, 4_X_3, 3_X_1, and 1_X_2 in the above example (where "X" denoted the middle stimulus, regardless of whether it was a random or fixed pattern element) occurred with high probability, as they can be formed both by two sequence elements comprising a random one or two random ones comprising a sequence element, thus, their last elements could be predicted either from the fixed pattern or from the combination of pattern and random elements. In contrast, triplets like 1_X_3 or 4_X_1 were considered low-probability, as they could only occur when both the first and last elements were random. In total, there were 16 possible high-probability triplets and 48 low-probability triplets.

As a result, the last elements of high-probability triplets were more predictable, and thus responses to them tended to be faster and more accurate. Each high-probability triplet appeared approximately five times more frequently than each low-probability one. Specifically, high-probability triplets occurred with a ∼4% likelihood per trial and 62.5% (16 × 4%) per block, while low-probability triplets appeared with ∼0.8% per trial and 37.5% (48 × 0.8%) per block (see Figure 7). The difference in reaction time (RT) and/or accuracy between the last elements of high- and low-probability triplets was taken as a behavioural index of statistical learning. For analytic purposes, data were segmented into epochs, each consisting of 5 blocks, that is, a total of 85 × 5 trials.

**Figure 7.**
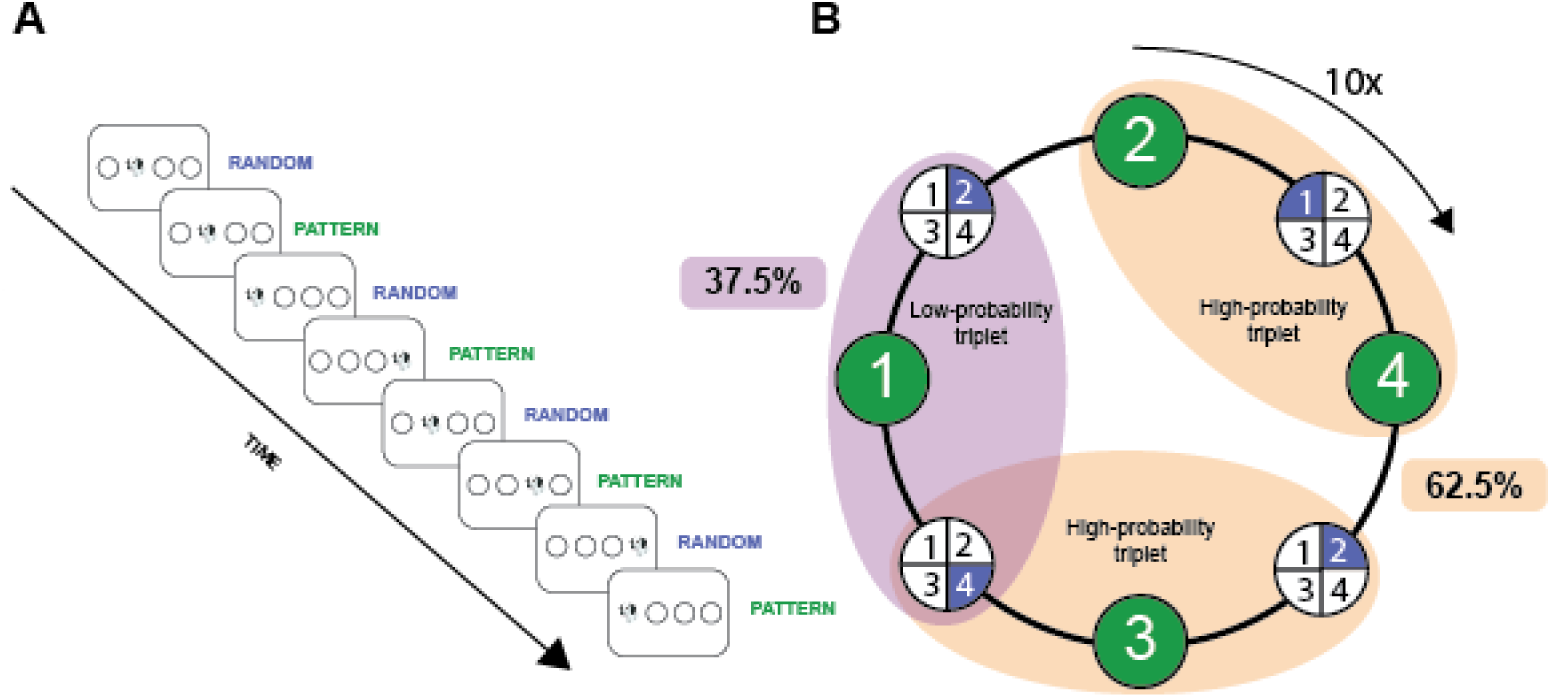
Structure of the Alternating Serial Reaction Time Task (ASRT) A. Participants were instructed to press the button corresponding to the location of the stimulus (a dog’s head). The order of stimuli followed a hidden alternating pattern: every first element was randomly selected (blue), appearing in any of the four circles, and every second element followed a fixed sequence (green), referred to as the pattern element. This resulted in an eight-element alternating sequence that was repeated 10 times within each block. B. Due to this alternating structure, certain three-element combinations of stimuli (called triplets) occurred with different probabilities. High-probability triplets (orange) can be formed when the third stimulus follows the hidden pattern, and may start with either a random or a pattern element. These make up 62.5% of all triplet occurrences. In contrast, low-probability triplets (purple) can only occur in random-pattern-random configurations and thus represent 37.5% of the total.

#### Paired Associate Learning Task

The Paired Associate Learning Task (PALT), developed for the Hungarian language by Nagy et al.^67^, is a computerized paradigm designed to assess declarative memory and is based on the revised version of a task by Cohn & Moscovitch^68^. During the task, two drawings – depicting either objects or animals – were simultaneously presented on the screen. The task consisted of several phases. In the initial learning phase, participants were asked to name each pair of images aloud as they appeared sequentially, without any instruction to memorize them. After naming the pictures, the experimenter advanced the display to the next image pair. A total of 23 image pairs were shown during this phase.

Subsequent retrieval phases involved the presentation of image pairs again, and participants were asked to indicate whether they had previously seen each item during the learning phase. Four possible types of image pairs were presented: old-old (both previously seen), new-new (none of the images were previously seen), old-new, and new-old. In cases where participants responded that both items had been presented before, participants were further asked to determine whether the pair had originally appeared together (“original” order) in the learning phase or had been recombined (“rearranged” order). Although the images did not repeat within the retrieval phase, participants were specifically instructed to give their responses solely based on the learning phase. Responses were recorded by the experimenter to avoid confusion due to the complexity of response options, and therefore, reaction time was not measured.

Unlike the original implementation by Nagy et al.^67^, in our study, participants were not informed in the learning phase that they would later be tested on the retrieval of the images following the naming task. As such, they did not expect a memory test, allowing for an incidental learning design. Additionally, in contrast to the original continuous task format, we divided the task into discrete retrieval blocks: three blocks during the first day and one block during the second day. Each block included eight unique image pairs, with no repetition across blocks to prevent interference with consolidation processes.

### Stimulation protocol

rTMS was administered using a Deymed DuoMAG XT stimulator over the DLPFC, targeting Brodmann area 9, corresponding to the F3 and F4 positions of the international 10–20 EEG system, which we located using a standard EEG cap (which has been shown to be a reasonably reliable method of neuronavigation^64^). Motor thresholding was individually established for each participant, by single TMS pulses delivered over the participants’ right primary motor cortex (M1) as initially localized by the C4 electrode of a 10-20 system EEG cap. The C4 electrode location was used as an initial anatomical reference point. From this location, we conducted a systematic searching procedure with small coil displacements to identify each individual’s motor hotspot for the right first dorsal interosseous (FDI) muscle. The hotspot was defined as the scalp location at which TMS elicited the largest and most reliable EMG responses in the resting FDI muscle. Resting motor threshold was then determined over the individually identified hotspot by gradually increasing stimulation intensity until minimum stimulation intensity that elicited visible muscle twitches in the hand or evoked motor potentials (MEPs) with an amplitude of at least 100 μV. This approach ensured that motor thresholding was based on functionally localized motor representations rather than a fixed scalp location. Stimulation on the first day was applied at 100% of each participant’s individual resting motor threshold in the active stimulation groups. A 1 Hz inhibitory stimulation protocol was used, delivering 300 pulses in each stimulation run, over 5 minutes to either the right or left hemisphere in the active right and active left groups, respectively. In the active bilateral group, stimulation was applied consecutively to each hemisphere for 2.5-2.5 minutes. In the sham condition, the stimulation coil was held perpendicular to the scalp (following the configuration described by^9^), preventing effective stimulation. However, the coil was still positioned on the participant’s head, allowing them to believe they were receiving stimulation. This approach was used to control for potential placebo effects.

### Procedure

Participants completed an online screening questionnaire designed to exclude individuals with health-related conditions (e.g., neurological or psychiatric disorders) that would preclude participation. Only those meeting the inclusion criteria were enrolled in the study, which consisted of two experimental days held 24 hours apart. Prior to the first day, participants were randomly assigned to one of the groups (active left, right, bilateral DLPFC stimulation, or sham stimulation) and a counterbalanced task order, determining whether they would first complete the ASRT or the PALT task. This assignment remained fixed throughout the entire experiment for each participant.

On Day 1, after providing written informed consent, the location of Brodmann area 9 and the M1 were identified using a 10-20 system EEG cap, followed by the assessment of each participant’s resting motor threshold. Participants then completed a familiarization phase of the ASRT task (two blocks of stimuli without embedded sequences) and the learning phase of the PALT task, with task order determined by prior randomization. Participants received a 5-minute rTMS run targeting either the right DLPFC, left DLPFC, or sham stimulation. In the bilateral stimulation group, 2.5 minutes of stimulation was delivered to each hemisphere consecutively - the order of the hemispheres was also randomized across participants. Immediately thereafter, participants completed the first epoch of the sequence-containing ASRT task (comprising five blocks, approximately five minutes in total) and the first three retrieval phases of the PALT task in a randomized order. This rTMS–ASRT–PALT cycle was repeated five times, with the order of the ASRT and PALT tasks alternating based on the assigned condition. During the second and fourth cycles, the PALT task was replaced with short breaks. In the second day, held 24 hours later, participants again completed one epoch of the ASRT task and a retrieval phase of the PALT tasks in the same randomized order as on the first day. The experimental procedure is illustrated in Figure 8. To ensure that baseline cognitive differences between groups did not confound the results (e.g., one group performing better due to inherently higher cognitive capacity), we administered the CSPAN^69^, measuring complex working memory tasks to detect any such pre-existing group-level advantages (see Table 2).

**Figure 8.**
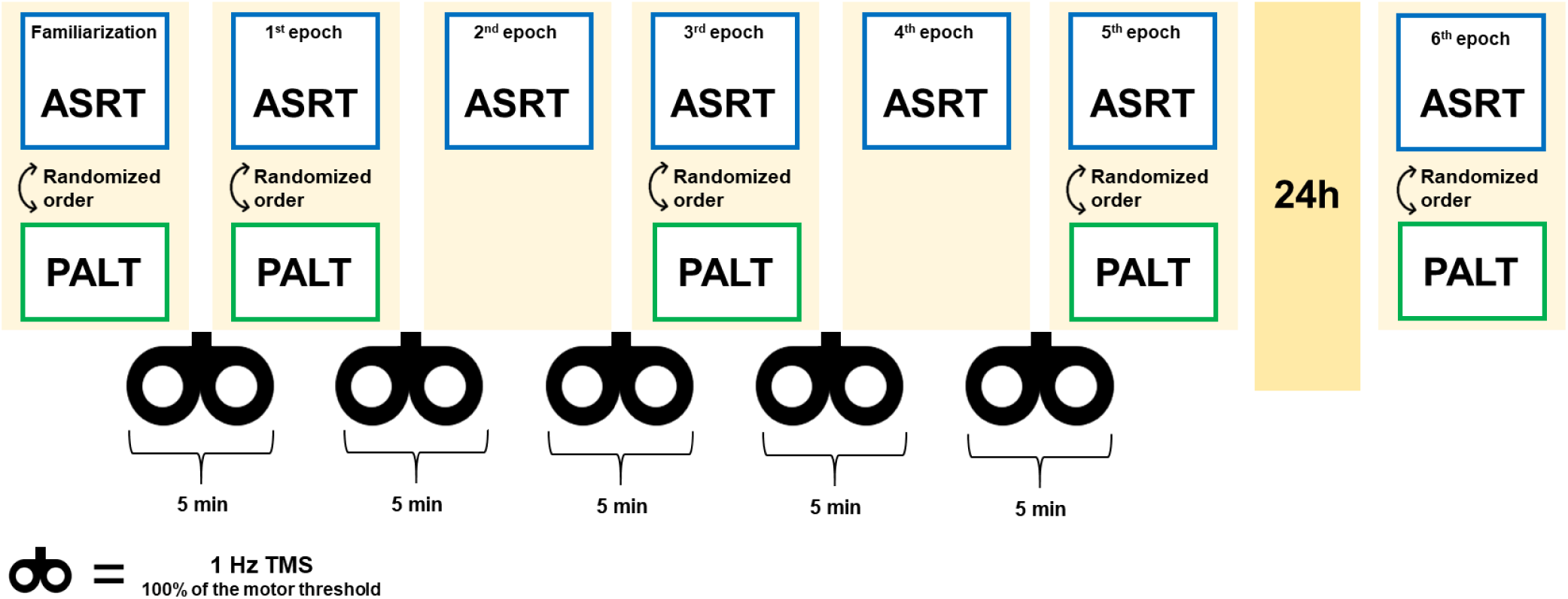
Study design. The experiment consisted of two sessions separated by a 24-hour delay. At the beginning of the first session, participants were randomly assigned to one of four stimulation groups (sham, right DLPFC, left DLPFC, bilateral DLPFC). Task order was also randomized: each participant was assigned to begin the session with either the ASRT task or the PALT task, and this order was kept consistent throughout all subsequent blocks. Following this, participants completed a familiarization phase of the ASRT task (which was two blocks of randomized trials, following no sequence) and the learning phase of the PALT task. These were followed by the first rTMS stimulation (1 Hz, 5 minutes, at 100% of the participant’s individual motor threshold). Stimulation was applied to the assigned site: for bilateral stimulation, 2.5 minutes were applied to each hemisphere. After the first stimulation, five blocks of task performance alternated with five stimulation runs. Notably, only the ASRT task was administered following the second and fourth stimulation blocks. In total, participants completed five task–stimulation cycles during the first day only. After a 24-hour delay, participants returned for the second session, during which they completed a sixth epoch of the ASRT task and the retrieval phase of the PALT task without stimulation.

### Statistical analysis

All statistical analyses were conducted in RStudio (Version 2024.12.0) or JASP (Version 0.19.3)^70,71^. Linear mixed-effects models were fitted using the *mixed* function from the *afex* package^72^. We used the default method to approximate degrees of freedom for fixed effects in all models. Post-hoc contrasts were computed using the *emmeans* package^73^. All figures and data preprocessing were conducted in Python (Version 3.13) using the *pandas*^74^, *numpy*^75^, *os*, *io*^76^, *seaborn*^77^, and *matplotlib*^78^ libraries. In all analyses, we determined an alpha level of 0.05.

#### Alternating Serial Reaction Time Task

Trials were categorized into high- and low-probability in a sliding window manner based on whether they were the last element of a high- or a low-probability triplet (henceforth simply referred to as high- and low-probability triplets). We excluded the first five trials of each block as they were random and served as practice trials, and trials that were the last element of repetitions (e.g., 1-1-1, 2-2-2) or trills (e.g., 1-2-1, 2-3-2), as participants may show a pre-existing tendency of faster motor responses to such occurrences^43^. Furthermore, from the analyses of RT, inaccurate trials were excluded.

To investigate group differences in statistical learning on RTs, we conducted a linear mixed-effect model to analyze RTs as a continuous outcome variable and a generalized linear mixed model to analyze accuracy, with responses coded as 0 (incorrect) and 1 (correct). The fixed effects included Epoch (1–6), Triplet Type (high- vs. low-probability), Group (sham, right, left, and bilateral stimulation groups), and their interactions. The models allowed for participant-specific intercepts and correlated slopes for within-subject variables. We used a top-down approach in model selection, starting from the most complex model (allowing for participant-specific intercept and correlated slopes for the interaction of within-subject variables) and simplified it gradually until we got a converging model. The fixed effects of the generalized linear mixed model were tested using Type III Wald chi-square tests with the *anova()* function from the car package. In addition, full model coefficients, including all interaction terms, are reported using the *tab_model()* function from the *sjPlot* package^79^.

To test for the robustness of our results, we repeated the main analyses using Bayesian methods implemented by the *brm()* function of the brms package. For the RT, we fitted a Bayesian linear mixed model (estimated using Hamiltonian Monte Carlo sampling with 3 chains of 2000 iterations and a warmup of 500, using the brms default priors). We applied the same model structure as in the frequentist version (outcome variable: RT, fixed factors: Epoch, Triplet type, Group, random factor: Participant number, random intercept included). For the accuracy, we applied a Bayesian generalized linear mixed model using the same estimation parameters as for the RT. The model structure, again, matched the frequentist version of the analysis (outcome variable: accuracy, fixed factors: Epoch, Triplet type, Group, random factor: Participant number, random intercept included).

#### Paired Associate Learning Task

To compare the groups in episodic retrieval performance, we conducted mixed-design analyses of variance (ANOVAs), as all linear mixed-effects models failed to converge on these data. We used Epoch (four levels) as the within-subject factor and Group (sham, right, left, and bilateral stimulation groups) as the between-subject factor. We ran three ANOVAs, as three scores were derived from participants’ responses. Item memory was calculated by subtracting the false alarm rate (incorrect Old– Old responses to New–New pairs) from the hit rate for rearranged Old–Old pairs. The association learning index was defined as the difference between the hit rate for original and rearranged Old–Old pairs. The recollection index was computed by subtracting the false alarm rate (incorrect original responses to rearranged Old–Old pairs) from the hit rate for original Old–Old pairs. Prior to the analysis, we tested whether the assumptions of the ANOVA were met. Specifically, the assumption of sphericity was assessed using Mauchly’s test, and normality was evaluated using Q–Q plots. When the assumption of sphericity was violated, the Greenhouse–Geisser correction was applied to adjust the degrees of freedom. To test null effects, we applied Bayesian mixed-design ANOVAs on the PALT scores. The Bayes Factor_10_ (BF_10_) was calculated using JASP to compare the null and alternative hypotheses. Null models included the subject variable and random slopes. Based on the BF10 values, we calculated BF_incl_ values across matched models. BF_incl_ values indicate the likeliness of a model that does include the given effect vs. the one that does not. The BF_incl_ values above one support the inclusion of the factor in the model, while values below one support the exclusion^80^. These values can be interpreted similarly to the BF_10_ scores: following the interpretation guidelines by Wagenmakers et al.^81^, values around 1 indicate no substantial evidence for either hypothesis. BF_incl_ values between 1 and 3 indicate anecdotal evidence for H₁, while values between 3 and 10 suggest moderate evidence. Conversely, values between 0.33 and 1 indicate anecdotal evidence for H₀, and values between 0.1 and 0.33 suggest moderate support for H₀. A BF_incl_ below 0.1 is considered strong evidence in favor of the null hypothesis, indicating that the groups likely do not differ.

Lastly, we performed some exploratory analysis that may help us understand our results more. First, as an indicator of exploratory behaviour in the task, we tested the RT variability on the ASRT task by conducting a Levene’s test, using Group as the independent variable. Post hoc analyses were corrected for multiple comparisons using the Holm method. In addition, as working memory is a relevant factor in the DLPFC-statistical learning relationship^21^, we investigated whether working memory capacity is related to statistical learning performance. We ran a correlation between total learning score (calculated as the difference between low- and high-probability triplets across all sequential epochs) and baseline counting span scores, using Pearson’s correlation. Please note that these analyses were exploratory and not part of our predetermined analysis plan, and they serve to further explain the results of the main analyses; for justification, see the Discussion section.

## Acknowledgments

This work was supported by the French National Grant Agency (ANR-22-CPJ1-0042-01 and ANR-24-CE37-5807); the National Brain Research Program project NAP2022-I-2/2022; and the Hungarian Scientific Research Fund (NKFIH 153150) (all awarded to D.N.). Supported by the EKÖP-25 University Excellence Scholarship Program of the Ministry for Culture and Innovation from the source of the National Research, Development, and Innovation Fund (EKÖP-25-3-I-ELTE-349, to ZVP). The Authors thank Dóra Osztényi, Noémi Papp, and Ádám Bicski for their work in the data collection.

## Data availability statement

Data and codes for analysis are available on the following link: https://osf.io/e2rzs, DOI 10.17605/OSF.IO/E2RZS

## Declaration of Generative AI and AI-assisted technologies in the writing process

During the preparation of this work the authors used ChatGPT (GPT-5 models) in order to correct grammatical and stylistic mistakes, and to add documentation to the analysis code files. After using this tool/service, the authors reviewed and edited the content as needed and take full responsibility for the content of the publication.

## References

1. Poldrack, R. A. & Packard, M. G. Competition among multiple memory systems: converging evidence from animal and human brain studies. Neuropsychologia 41, 245–251 (2003).

2. Borragán, G., Slama, H., Destrebecqz, A. & Peigneux, P. Cognitive Fatigue Facilitates Procedural Sequence Learning. Front. Hum. Neurosci. 10, (2016).

3. Zhong, L. et al. Unsupervised pretraining in biological neural networks. Nature 644, 741–748 (2025).

4. Thompson-Schill, S. L., Ramscar, M. & Chrysikou, E. G. Cognition Without Control: When a Little Frontal Lobe Goes a Long Way. Curr Dir Psychol Sci 18, 259–263 (2009).

5. Sherman, B. E. & Turk-Browne, N. B. Statistical prediction of the future impairs episodic encoding of the present. Proc. Natl. Acad. Sci. U.S.A. 117, 22760–22770 (2020).

6. Daw, N. D., Niv, Y. & Dayan, P. Uncertainty-based competition between prefrontal and dorsolateral striatal systems for behavioral control. Nature Neuroscience 8, 1704–1711 (2005).

7. Virag, M. et al. Competition between frontal lobe functions and implicit sequence learning: evidence from the long-term effects of alcohol. Exp Brain Res 233, 2081–2089 (2015).

8. Pedraza, F. et al. Evidence for a competitive relationship between executive functions and statistical learning. *npj Sci*. Learn. 9, 30 (2024).

9. Ambrus, G. G. et al. When less is more: Enhanced statistical learning of non-adjacent dependencies after disruption of bilateral DLPFC. Journal of Memory and Language 114, 104144 (2020).

10. Tóth-Fáber, E., Nemeth, D. & Janacsek, K. Lifespan developmental invariance in memory consolidation: evidence from procedural memory. PNAS Nexus 2, pgad037 (2023).

11. Nemeth, D., Janacsek, K. & Fiser, J. Age-dependent and coordinated shift in performance between implicit and explicit skill learning. Front. Comput. Neurosci. 7, (2013).

12. Juhasz, D., Nemeth, D. & Janacsek, K. Is there more room to improve? The lifespan trajectory of procedural learning and its relationship to the between- and within-group differences in average response times. PLoS ONE 14, e0215116 (2019).

13. Nemeth, D., Janacsek, K., Polner, B. & Kovacs, Z. A. Boosting Human Learning by Hypnosis. Cerebral Cortex 23, 801–805 (2013).

14. Smalle, E. H. M., Muylle, M., Duyck, W. & Szmalec, A. Less is more: Depleting cognitive resources enhances language learning abilities in adults. Journal of Experimental Psychology: General 150, 2423–2434 (2021).

15. Smalle, E. H. M., Daikoku, T., Szmalec, A., Duyck, W. & Möttönen, R. Unlocking adults’ implicit statistical learning by cognitive depletion. Proc. Natl. Acad. Sci. U.S.A. 119, e2026011119 (2022).

16. Szücs-Bencze, L. et al. Enhancing retrieval capacity of the predictive brain through dorsolateral prefrontal cortex intervention. Cerebral Cortex 35, bhaf005 (2025).

17. Janacsek, K., Ambrus, G. G., Paulus, W., Antal, A. & Nemeth, D. Right Hemisphere Advantage in Statistical Learning: Evidence From a Probabilistic Sequence Learning Task. Brain Stimulation 8, 277–282 (2015).

18. Roser, M. E., Fiser, J., Aslin, R. N. & Gazzaniga, M. S. Right Hemisphere Dominance in Visual Statistical Learning. Journal of Cognitive Neuroscience 23, 1088–1099 (2010).

19. Wolford, G., Miller, M. B. & Gazzaniga, M. The left hemisphere’s role in hypothesis formation. The Journal of Neuroscience 20, RC64 (2000).

20. Greeley, B. & Seidler, R. D. Differential effects of left and right prefrontal cortex anodal transcranial direct current stimulation during probabilistic sequence learning. Journal of Neurophysiology 121, 1906–1916 (2019).

21. Smittenaar, P., FitzGerald, T. H. B., Romei, V., Wright, N. D. & Dolan, R. J. Disruption of Dorsolateral Prefrontal Cortex Decreases Model-Based in Favor of Model-free Control in Humans. Neuron 80, 914–919 (2013).

22. Galea, J. M., Albert, N. B., Ditye, T. & Miall, R. C. Disruption of the Dorsolateral Prefrontal Cortex Facilitates the Consolidation of Procedural Skills. Journal of Cognitive Neuroscience 22, 1158–1164 (2009).

23. Savic, B., Cazzoli, D., Müri, R. & Meier, B. No effects of transcranial DLPFC stimulation on implicit task sequence learning and consolidation. Sci Rep 7, 9649 (2017).

24. Savic, B., Müri, R. & Meier, B. High Definition Transcranial Direct Current Stimulation Does Not Modulate Implicit Task Sequence Learning and Consolidation. Neuroscience 414, 77–87 (2019).

25. Balconi, M. Dorsolateral prefrontal cortex, working memory and episodic memory processes: insight through transcranial magnetic stimulation techniques. Neurosci. Bull. 29, 381–389 (2013).

26. Ruchkin, D. S., Grafman, J., Cameron, K. & Berndt, R. S. Working memory retention systems: A state of activated long-term memory. Behav Brain Sci 26, 709–728 (2003).

27. Habib, R., Nyberg, L. & Tulving, E. Hemispheric asymmetries of memory: the HERA model revisited. Trends in Cognitive Sciences 7, 241–245 (2003).

28. Tulving, E., Kapur, S., Craik, F. I., Moscovitch, M. & Houle, S. Hemispheric encoding/retrieval asymmetry in episodic memory: positron emission tomography findings. PNAS 91, 2016–2020 (1994).

29. Smalle, E. H. M., Panouilleres, M., Szmalec, A. & Möttönen, R. Language learning in the adult brain: disrupting the dorsolateral prefrontal cortex facilitates word-form learning. Sci Rep 7, 13966 (2017).

30. Filoteo, J. V., Lauritzen, S. & Maddox, W. T. Removing the Frontal Lobes: The Effects of Engaging Executive Functions on Perceptual Category Learning. Psychol Sci 21, 415–423 (2010).

31. Tóth, B. et al. Dynamics of EEG functional connectivity during statistical learning. Neurobiology of Learning and Memory 144, 216–229 (2017).

32. Park, J., Janacsek, K., Nemeth, D. & Jeon, H.-A. Reduced functional connectivity supports statistical learning of temporally distributed regularities. NeuroImage 260, 119459 (2022).

33. Zwart, F. S., Vissers, C. Th. W. M., Kessels, R. P. C. & Maes, J. H. R. Procedural learning across the lifespan: A systematic review with implications for atypical development. Journal of Neuropsychology 13, 149–182 (2019).

34. Smith, E. E. et al. Spatial versus Object Working Memory: PET Investigations. Journal of Cognitive Neuroscience 7, 337–356 (1995).

35. Wager, T., Insler, R. & Smith, E. The neural bases of distracter-resistant working memory. Nat Prec https://doi.org/10.1038/npre.2011.5526.1 (2011) doi:10.1038/npre.2011.5526.1.

36. Frank, S. M. et al. Fundamental Differences in Visual Perceptual Learning between Children and Adults. Current Biology 31, 427–432.e5 (2021).

37. Divjak, D. & Milin, P. Exploring and Exploiting Uncertainty: Statistical Learning Ability Affects How We Learn to Process Language Along Multiple Dimensions of Experience. Cognitive Science 44, e12835 (2020).

38. Pesthy, O. et al. Intact predictive processing in autistic adults: evidence from statistical learning. Sci Rep 13, 11873 (2023).

39. Wilson, R. C., Bonawitz, E., Costa, V. D. & Ebitz, R. B. Balancing exploration and exploitation with information and randomization. Current Opinion in Behavioral Sciences 38, 49–56 (2021).

40. Feng, S. F., Wang, S., Zarnescu, S. & Wilson, R. C. The dynamics of explore–exploit decisions reveal a signal-to-noise mechanism for random exploration. Sci Rep 11, 3077 (2021).

41. Wu, C. M., Schulz, E., Pleskac, T. J. & Speekenbrink, M. Time pressure changes how people explore and respond to uncertainty. Sci Rep 12, 4122 (2022).

42. Oberauer, K. Working Memory and Attention – A Conceptual Analysis and Review. Journal of Cognition 2, 36 (2019).

43. Olivers, C. N., Meijer, F. & Theeuwes, J. Feature-based memory-driven attentional capture: visual working memory content affects visual attention. Journal of experimental psychology: human perception and performance 32, 1243 (2006).

44. Jonides, J. et al. Spatial working memory in humans as revealed by PET. Nature 363, 623–625 (1993).

45. Dickey, C. W. et al. Transcranial magnetic stimulation to the dorsolateral prefrontal cortex modulates single-neuron activity in humans. Preprint at 10.64898/2026.03.15.711839 (2026).

46. Fried, P. J., Rushmore, R. J., Moss, M. B., Valero-Cabré, A. & Pascual-Leone, A. Causal evidence supporting functional dissociation of verbal and spatial working memory in the human dorsolateral prefrontal cortex. Eur J of Neuroscience 39, 1973–1981 (2014).

47. Lycke, C., Specht, K., Ersland, L. & Hugdahl, K. An fMRI study of phonological and spatial working memory using identical stimuli. Scandinavian J Psychology 49, 393–401 (2008).

48. Reuter-Lorenz, P. A. et al. Age Differences in the Frontal Lateralization of Verbal and Spatial Working Memory Revealed by PET. Journal of Cognitive Neuroscience 12, 174–187 (2000).

49. Smith, E. E., Jonides, J. & Koeppe, R. A. Dissociating Verbal and Spatial Working Memory Using PET. Cerebral Cortex 6, 11–20 (1996).

50. Wan, Q. & Sloutsky, V. M. Working memory shapes information sampling and attention allocation across development. Journal of Experimental Psychology: General 155, 479–498 (2026).

51. Hayden, B. Y., Heilbronner, S. R. & Yoo, S. B. M. Rethinking the centrality of brain areas in understanding functional organization. Nat Neurosci 29, 267–278 (2026).

52. Rosen, M. C. & Freedman, D. J. How distributed is the brain-wide network that is recruited for cognition? Nat. Rev. Neurosci. 27, 138–150 (2026).

53. Sandrini, M., Cappa, S. F., Rossi, S., Rossini, P. M. & Miniussi, C. The Role of Prefrontal Cortex in Verbal Episodic Memory: rTMS Evidence. Journal of Cognitive Neuroscience 15, 855–861 (2003).

54. Arabacı, G., Cakir, B. S. & Parris, B. A. The effect of high-frequency rTMS over left DLPFC and fluid abilities on goal neglect. Brain Struct Funct 229, 1073–1086 (2024).

55. Cui, H. et al. A novel dual-site OFC-dlPFC accelerated repetitive transcranial magnetic stimulation for depression: a pilot randomized controlled study. Psychol. Med. 54, 3849–3862 (2024).

56. Izuma, K. et al. A Causal Role for Posterior Medial Frontal Cortex in Choice-Induced Preference Change. J. Neurosci. 35, 3598–3606 (2015).

57. Kim, L., Chun, M. H., Kim, B. R. & Lee, S. J. Effect of Repetitive Transcranial Magnetic Stimulation on Patients with Brain Injury and Dysphagia. Ann Rehabil Med 35, 765 (2011).

58. Kurkin, S. et al. Transcranial Magnetic Stimulation of the Dorsolateral Prefrontal Cortex Increases Posterior Theta Rhythm and Reduces Latency of Motor Imagery. Sensors 23, 4661 (2023).

59. Schluter, R. S., Van Holst, R. J. & Goudriaan, A. E. Effects of Ten Sessions of High Frequency Repetitive Transcranial Magnetic Stimulation (HF-rTMS) Add-on Treatment on Impulsivity in Alcohol Use Disorder. Front. Neurosci. 13, 1257 (2019).

60. Shang, Y. et al. Theta-burst transcranial magnetic stimulation induced functional connectivity changes between dorsolateral prefrontal cortex and default-mode-network. Brain Imaging and Behavior 14, 1955–1963 (2020).

61. Sun, T. & Yu, Q. Effects of repetitive transcranial magnetic stimulation on upper extremity motor function in stroke survivors: study protocol of a randomized sham-controlled trial. Front. Neurol. 16, 1669862 (2026).

62. Vanderhasselt, M.-A., De Raedt, R., Baeken, C., Leyman, L. & D’haenen, H. The influence of rTMS over the left dorsolateral prefrontal cortex on Stroop task performance. Exp Brain Res 169, 279–282 (2006).

63. Caulfield, K. A. et al. Neuronavigation maximizes accuracy and precision in TMS positioning: Evidence from 11,230 distance, angle, and electric field modeling measurements. Brain Stimulation 15, 1192–1205 (2022).

64. Herwig, U., Satrapi, P. & Schönfeldt-Lecuona, C. Using the International 10-20 EEG System for Positioning of Transcranial Magnetic Stimulation. Brain Topogr 16, 95–99 (2003).

65. Hebel, T. et al. A direct comparison of neuronavigated and non-neuronavigated intermittent theta burst stimulation in the treatment of depression. Brain Stimulation 14, 335–343 (2021).

66. Howard, J. H. & Howard, D. V. Age differences in implicit learning of higher order dependencies in serial patterns. Psychology and Aging 12, 634–656 (1997).

67. Nagy M., Kónya A. & Király I. Az Asszociatív Emlékezet Fejlődése 6–10 Éves Kor És Fiatal Felnőttkor Között. Pszichológia 33, 169–184 (2013).

68. Cohn, M. & Moscovitch, M. Dissociating measures of associative memory: Evidence and theoretical implications. Journal of Memory and Language 57, 437–454 (2007).

69. Case, R., Kurland, D. M. & Goldberg, J. Operational efficiency and the growth of short-term memory span. Journal of Experimental Child Psychology 33, 386–404 (1982).

70. JASP Team. JASP (Version 0.18.3)[Computer software]. (2024).

71. RStudio: Integrated Development Environment for R. RStudio Team (2015).

72. Singmann, H. et al. afex: Analysis of Factorial Experiments. (2024).

73. Lenth, R. V. Emmeans: Estimated Marginal Means, Aka Least-Squares Means. (2024).

74. Reback, J., et al. pandas-dev/pandas: Pandas 1.0.3. Zenodo 10.5281/ZENODO.3715232 (2020).

75. Harris, C. R. et al. Array programming with NumPy. Nature 585, 357–362 (2020).

76. David J. H. Shih. io: A Unified Framework for Input-Output Operations in R. 0.3.2 10.32614/CRAN.package.io (2014).

77. Waskom, M. et al. mwaskom/seaborn: v0.8.1 (September 2017). Zenodo 10.5281/ZENODO.883859 (2017).

78. Hunter, J. D. Matplotlib: A 2D Graphics Environment. Comput. Sci. Eng. 9, 90–95 (2007).

79. Lüdecke, D. sjPlot: Data Visualization for Statistics in Social Science. 2.9.0 10.32614/CRAN.package.sjPlot (2013).

80. Peter Rosenfeld, J. & Olson, J. M. Bayesian Data Analysis: A Fresh Approach to Power Issues and Null Hypothesis Interpretation. Appl Psychophysiol Biofeedback 46, 135–140 (2021).

81. Wagenmakers, E.-J., Wetzels, R., Borsboom, D. & Van Der Maas, H. L. J. Why psychologists must change the way they analyze their data: The case of psi: Comment on Bem (2011). Journal of Personality and Social Psychology 100, 426–432 (2011).

82. Hou, M., de Chastelaine, M., and Rugg, M.D. Age differences in the neural correlates of recollection: transient versus sustained fMRI effects. Neurobiol. Aging 131, 132–143 (2023). 10.1016/J.NEUROBIOLAGING.2023.07.001.

83. Vilberg, K.L., and Rugg, M.D. Temporal dissociations within the core recollection network. Cogn. Neurosci. 5, 77–84 (2014). 10.1080/17588928.2013.860088.

84. Hockley, W.E., and Cristi, C. Tests of encoding tradeoffs between item and associative information. Mem. Cognit. 24, 202–216 (1996). 10.3758/BF03200881/METRICS.

