## Supplementary Materials for "Inhibiting the right dorsolateral prefrontal cortex selectively enhances unsupervised statistical learning"

**Supplementary Materials for the manuscript "Inhibiting the right dorsolateral prefrontal cortex selectively enhances unsupervised statistical learning"**

Orsolya Pesthy*[1,2], Zsuzsanna Viktória Pesthy*[2,3,4], Teodóra Vékony[5], Karolina Janacsek[4,6], Dániel Fabó[7,8,9], Dezso Nemeth[1,5,10]

*1 Centre de Recherche en Neurosciences de Lyon, INSERM, CNRS, Université Claude Bernard Lyon 1, CRNL U1028 UMR5292, Bron, France*

*2 Brain, Memory and Language Research Group, Institute of Cognitive Neuroscience and Psychology, HUN-REN Research Centre for Natural Sciences, Budapest, Hungary*

*3 Doctoral School of Psychology, ELTE Eötvös Loránd University, Budapest, Hungary*

*4 Institute of Psychology, ELTE Eötvös Loránd University, Budapest, Hungary*

*5 Gran Canaria Cognitive Research Center, Atlántico Medio University, Las Palmas de Gran Canaria, Spain*

*6 Centre for Thinking and Learning, Institute for Lifecourse Development, School of Human Sciences, Faculty of Education, Health and Human Sciences, University of Greenwich, London, United Kingdom*

*7 Flash Clinic, Budapest, Hungary*

*8 Department of Neurology, University of Szeged, Szeged, Hungary*

*9 Dept of Voice, Speech and Swallowing, Semmelweis University, Budapest, Hungary*

*10 BML-NAP Research Group, Institute of Psychology, Eötvös Loránd University & Institute of Cognitive Neuroscience and Psychology, HUN-REN Research Centre for Natural Sciences, Budapest, Hungary*

**These authors contributed equally to this article*

### Supplementary Results

#### Frequentist analyses: main analyses

**Table S1.** Full results of the mixed-effects models predicting reaction times (RT)

| | RT | | |
|---|---|---|---|
| **Predictors** | **Estimate** | **SE** | **95% CI** |
| (Intercept) | 356.88 | 3.32 | 350.38 – 363.39 |
| Right | -6.02 | 5.58 | -16.95 – 4.90 |
| Left | 3.72 | 5.89 | -7.83 – 15.27 |
| Bilateral | -4.39 | 5.81 | -15.77 – 6.99 |
| Epoch 1 | 20.79 | 0.45 | 19.92 – 21.67 |
| Epoch 2 | 8.05 | 0.45 | 7.17 – 8.93 |
| Epoch 3 | 0.65 | 0.45 | -0.23 – 1.53 |
| Epoch 4 | -3.91 | 0.45 | -4.79 – -3.02 |
| Epoch 5 | -7.97 | 0.45 | -8.86 – -7.08 |
| Triplet Type | -4.46 | 0.20 | -4.85 – -4.06 |
| Right × Epoch 1 | 0.77 | 0.75 | -0.70 – 2.25 |
| Left × Epoch 1 | 3.13 | 0.79 | 1.58 – 4.68 |
| Bilateral × Epoch 1 | -1.25 | 0.79 | -2.79 – 0.29 |
| Right × Epoch 2 | -1.41 | 0.76 | -2.89 – 0.08 |

| | | | |
|---|---|---|---|
| Left × Epoch 2 | 1.12 | 0.80 | -0.45 – 2.68 |
| Bilateral × Epoch 2 | 0.24 | 0.79 | -1.31 – 1.78 |
| Right × Epoch 3 | -1.35 | 0.76 | -2.84 – 0.13 |
| Left × Epoch 3 | 0.64 | 0.80 | -0.92 – 2.20 |
| Bilateral × Epoch 3 | -0.65 | 0.79 | -2.20 – 0.90 |
| Right × Epoch 4 | -0.90 | 0.76 | -2.40 – 0.60 |
| Left × Epoch 4 | -3.07 | 0.80 | -4.63 – -1.50 |
| Bilateral × Epoch 4 | 2.08 | 0.79 | 0.53 – 3.64 |
| Right × Epoch 5 | 0.88 | 0.76 | -0.61 – 2.38 |
| Left × Epoch 5 | 0.78 | 0.80 | -0.79 – 2.35 |
| Bilateral × Epoch 5 | -0.28 | 0.79 | -1.83 – 1.28 |
| Right × Triplet Type | 0.83 | 0.34 | 0.16 – 1.49 |
| Left × Triplet Type | -0.78 | 0.36 | -1.47 – -0.08 |
| Bilateral × Triplet Type | 0.48 | 0.35 | -0.21 – 1.17 |
| Epoch 1 × Triplet Type | 2.79 | 0.45 | 1.92 – 3.67 |
| Epoch 2 × Triplet Type | 1.56 | 0.45 | 0.67 – 2.44 |
| Epoch 3 × Triplet Type | 0.75 | 0.45 | -0.13 – 1.64 |
| Epoch 4 × Triplet Type | -1.25 | 0.45 | -2.14 – -0.36 |

| | | | |
|---|---|---|---|
| Epoch 5 × Triplet Type | -2.82 | 0.45 | -3.71 – -1.93 |
| Right × Epoch 1 × Triplet Type | 0.03 | 0.75 | -1.45 – 1.50 |
| Left × Epoch 1 × Triplet Type | 0.44 | 0.79 | -1.11 – 1.99 |
| Bilateral × Epoch 1 × Triplet Type | -0.03 | 0.79 | -1.57 – 1.51 |
| Right × Epoch 2 × Triplet Type | -0.69 | 0.76 | -2.18 – 0.80 |
| Left × Epoch 2 × Triplet Type | 0.16 | 0.80 | -1.41 – 1.72 |
| Bilateral × Epoch 2 × Triplet Type | 0.11 | 0.79 | -1.43 – 1.65 |
| Right × Epoch 3 × Triplet Type | -0.54 | 0.76 | -2.03 – 0.94 |
| Left × Epoch 3 × Triplet Type | -0.42 | 0.80 | -1.98 – 1.14 |
| Bilateral × Epoch 3 × Triplet Type | 0.48 | 0.79 | -1.07 – 2.02 |
| Right × Epoch 4 × Triplet Type | 0.33 | 0.76 | -1.17 – 1.82 |
| Left × Epoch 4 × Triplet Type | 0.47 | 0.80 | -1.09 – 2.04 |
| Bilateral × Epoch 4 × Triplet Type | -1.09 | 0.79 | -2.64 – 0.46 |

| | | | |
|---|---|---|---|
| Right × Epoch 5 × Triplet Type | 0.84 | 0.76 | -0.66 – 2.33 |
| Left × Epoch 5 × Triplet Type | 0.46 | 0.80 | -1.11 – 2.04 |
| Bilateral × Epoch 5 × Triplet Type | -0.11 | 0.79 | -1.67 – 1.45 |
| **Random Effects** | | | |
| $\sigma^2$ | 6064.88 | | |
| $\tau_{00\ Subject}$ | 1039.40 | | |
| ICC | 0.15 | | |
| $N_{Subject}$ | 95 | | |
| Marginal $R^2$ / Conditional $R^2$ | 0.029 / 0.171 | | |

**Note.** Estimate = regression coefficient; SE = standard error; CI = 95% confidence interval. For Group, the sham stimulation group served as the reference category; for Epoch, Epoch 6 served as the reference category. $\sigma^2$ = residual variance; $\tau_{00}$ = random-intercept variance; ICC = intraclass correlation coefficient; Marginal $R^2$ represents the variance explained by the fixed effects, whereas Conditional $R^2$ represents the variance explained by both fixed and random effects.

**Table S2.** Full results of the mixed-effects models predicting accuracy (ACC).

| | ACC | | |
|---|---|---|---|
| **Predictors** | **Estimate** | **SE** | **95% CI** |
| (Intercept) | 19.74 | 1.38 | 17.20 – 22.65 |
| Right | 0.94 | 0.11 | 0.75 – 1.18 |
| Left | 1.29 | 0.16 | 1.02 – 1.64 |
| Bilateral | 0.81 | 0.10 | 0.64 – 1.03 |

| | | | |
|---|---|---|---|
| Epoch 1 | 1.26 | 0.03 | 1.19 – 1.32 |
| Epoch 2 | 1.08 | 0.03 | 1.03 – 1.13 |
| Epoch 3 | 1.02 | 0.02 | 0.97 – 1.07 |
| Epoch 4 | 0.95 | 0.02 | 0.91 – 1.00 |
| Epoch 5 | 0.91 | 0.02 | 0.87 – 0.96 |
| Triplet Type | 1.23 | 0.01 | 1.20 – 1.25 |
| Right × Epoch 1 | 1.05 | 0.05 | 0.97 – 1.14 |
| Left × Epoch 1 | 0.97 | 0.05 | 0.88 – 1.07 |
| Bilateral × Epoch 1 | 0.99 | 0.04 | 0.91 – 1.08 |
| Right × Epoch 2 | 0.94 | 0.04 | 0.87 – 1.02 |
| Left × Epoch 2 | 1.06 | 0.05 | 0.97 – 1.17 |
| Bilateral × Epoch 2 | 1.06 | 0.04 | 0.98 – 1.16 |
| Right × Epoch 3 | 1.03 | 0.04 | 0.95 – 1.11 |
| Left × Epoch 3 | 1.01 | 0.05 | 0.93 – 1.11 |
| Bilateral × Epoch 3 | 0.97 | 0.04 | 0.90 – 1.05 |
| Right × Epoch 4 | 0.97 | 0.04 | 0.90 – 1.05 |
| Left × Epoch 4 | 1.00 | 0.04 | 0.92 – 1.09 |
| Bilateral × Epoch 4 | 1.00 | 0.04 | 0.93 – 1.08 |

| | | | |
|---|---|---|---|
| Right × Epoch 5 | 1.00 | 0.04 | 0.93 – 1.08 |
| Left × Epoch 5 | 1.04 | 0.05 | 0.96 – 1.14 |
| Bilateral × Epoch 5 | 0.97 | 0.04 | 0.90 – 1.05 |
| Right × Triplet Type | 1.01 | 0.02 | 0.98 – 1.05 |
| Left × Triplet Type | 1.01 | 0.02 | 0.97 – 1.06 |
| Bilateral × Triplet Type | 1.00 | 0.02 | 0.97 – 1.04 |
| Epoch 1 × Triplet Type | 0.89 | 0.02 | 0.84 – 0.94 |
| Epoch 2 × Triplet Type | 0.98 | 0.02 | 0.93 – 1.03 |
| Epoch 3 × Triplet Type | 0.97 | 0.02 | 0.93 – 1.02 |
| Epoch 4 × Triplet Type | 1.04 | 0.02 | 0.99 – 1.09 |
| Epoch 5 × Triplet Type | 1.07 | 0.02 | 1.02 – 1.12 |
| Right × Epoch 1 × Triplet Type | 0.94 | 0.04 | 0.86 – 1.02 |
| Left × Epoch 1 × Triplet Type | 1.01 | 0.05 | 0.91 – 1.11 |
| Bilateral × Epoch 1 × Triplet Type | 1.03 | 0.04 | 0.95 – 1.12 |
| Right × Epoch 2 × Triplet Type | 1.03 | 0.04 | 0.95 – 1.11 |
| Left × Epoch 2 × Triplet Type | 1.06 | 0.05 | 0.97 – 1.17 |

| | | | |
|---|---|---|---|
| Bilateral × Epoch 2 × Triplet Type | 0.90 | 0.04 | 0.83 – 0.98 |
| Right × Epoch 3 × Triplet Type | 1.00 | 0.04 | 0.93 – 1.09 |
| Left × Epoch 3 × Triplet Type | 1.00 | 0.05 | 0.91 – 1.09 |
| Bilateral × Epoch 3 × Triplet Type | 1.06 | 0.04 | 0.98 – 1.14 |
| Right × Epoch 4 × Triplet Type | 0.99 | 0.04 | 0.92 – 1.06 |
| Left × Epoch 4 × Triplet Type | 0.99 | 0.04 | 0.90 – 1.08 |
| Bilateral × Epoch 4 × Triplet Type | 0.96 | 0.04 | 0.89 – 1.04 |
| Right × Epoch 5 × Triplet Type | 1.03 | 0.04 | 0.96 – 1.11 |
| Left × Epoch 5 × Triplet Type | 0.99 | 0.04 | 0.90 – 1.08 |
| Bilateral × Epoch 5 × Triplet Type | 1.01 | 0.04 | 0.94 – 1.09 |
| **Random Effects** | | | |
| $\sigma^2$ | 3.29 | | |
| $\tau_{00\ Subject}$ | 0.46 | | |
| ICC | 0.12 | | |
| $N_{Subject}$ | 95 | | |

| | |
|---|---|
| Observations | 194684 |
| Marginal $R^2$ / Conditional $R^2$ | 0.021 / 0.140 |

**Note.** Estimate = regression coefficient; SE = standard error; CI = 95% confidence interval. For Group, the sham stimulation group served as the reference category; for Epoch, Epoch 6 served as the reference category. $\sigma^2$ = residual variance; $\tau_{00}$ = random-intercept variance; ICC = intraclass correlation coefficient; Marginal R² represents the variance explaine

**Table S3.** Post hoc pairwise comparisons of the statistical learning effect on the RT, between stimulation groups (Group * Triplet type interaction).

| | Reaction time | | | |
|---|---|---|---|---|
| **Group comparison** | **Estimate of H-L difference** | **SE** | **95% CI** | **p** |
| Sham vs. Right | 3.20 | 1.14 | [0.98, 5.43] | .005 |
| Sham vs. Left | 0.70 | 1.13 | [−1.52, 2.91] | .538 |
| Sham vs. Bilateral | 2.71 | 1.12 | [0.53, 4.90] | .015 |
| Right vs. Left | −2.51 | 1.17 | [−4.80, −0.22] | .032 |
| Right vs. Bilateral | −0.49 | 1.16 | [−2.76, 1.78] | .671 |
| Left vs. Bilateral | 2.02 | 1.15 | [−0.23, 4.27] | .079 |

**Note**. Estimates represent between-group differences in the H–L reaction-time contrast (statistical learning effect), averaged across epochs. CI = confidence interval.

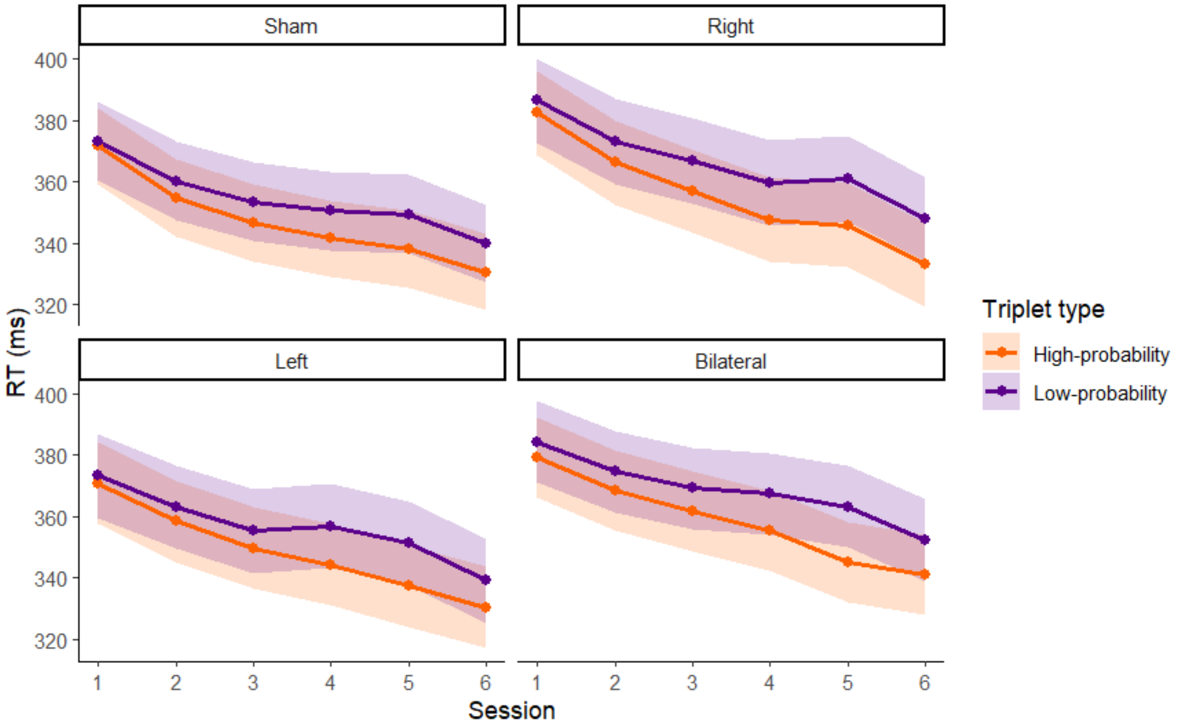


**Figure S1.** Frequentist model-estimated reaction times to high- and low-probability triplets across stimulation groups. The figure displays model-estimated marginal means of reaction times (in ms) for high-probability (orange) and low-probability (purple) triplets across six epochs (x-axis). Shaded areas represent 95% confidence intervals derived from the mixed-effects model. Panels correspond to the sham, right, left, and bilateral DLPFC stimulation groups. The difference between reaction times to high- and low-probability triplets reflects the magnitude of statistical learning over time.

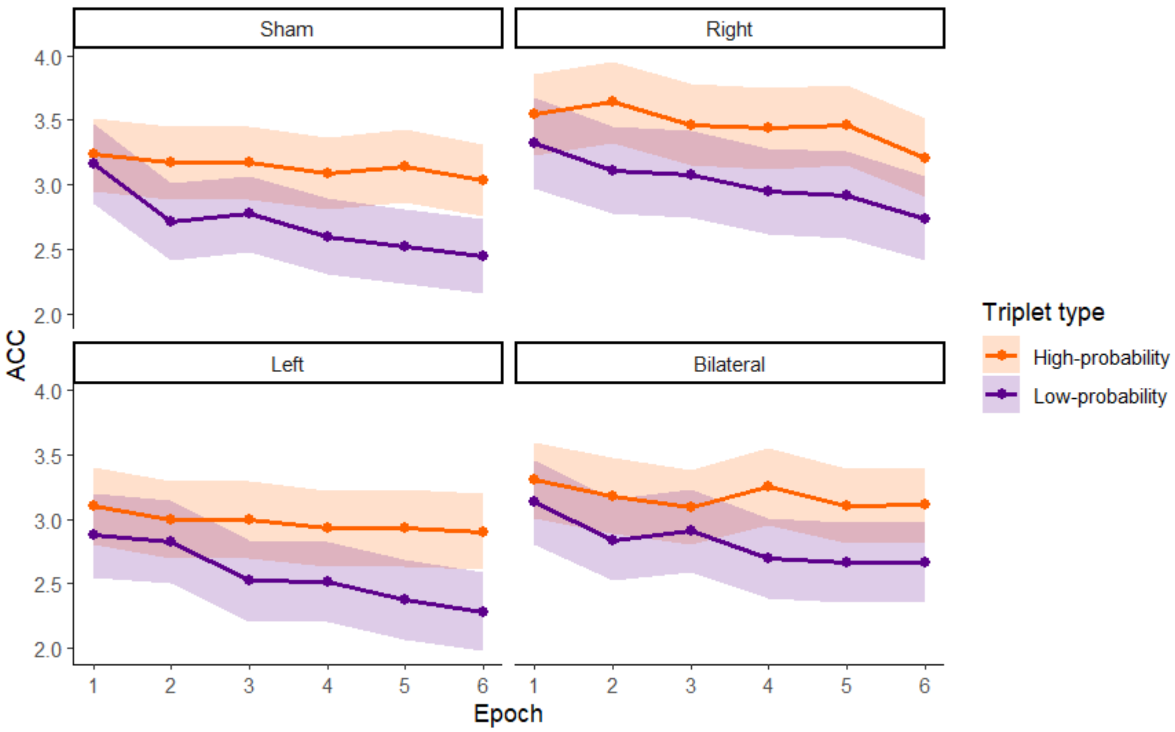


**Figure S2.** Frequentist model-estimated marginal means of accuracy for high-probability (orange) and low-probability (purple) triplets across six epochs (x-axis). Shaded areas represent 95% confidence intervals derived from the mixed-effects model. Panels correspond to the sham, right, left, and bilateral DLPFC stimulation groups. Differences in accuracy between high- and low-probability triplets reflect statistical learning effects across epochs. Additionally, the code used to generate the figures has been uploaded to OSF (TMS_plots_osf.ipynb) to facilitate transparency and reproducibility.

**Table S4.** Main effects and interactions of the generalized linear mixed-effect model on the accuracy of the Alternating Serial Reaction Time Task.

| **Effect** | **Chi-square ($\chi^2$)** | ***df*** | **p-value** | |
|---|---|---|---|---|
| Intercept | 1779.7748 | 1 | < 0.001 | *** |
| Group | 5.4579 | 3 | 0.141 | |
| Epoch | 141.5015 | 5 | < 0.001 | *** |
| Triplet Type (TT) | 363.9484 | 1 | < 0.001 | *** |
| Group × Epoch | 13.6893 | 15 | 0.549 | |

| | | | | |
|---|---|---|---|---|
| Group × TT | 2.5320 | 3 | 0.470 | |
| Epoch × TT | 32.3509 | 5 | < 0.001 | *** |
| Group × Epoch × TT | 14.4833 | 15 | 0.489 | |

#### Bayesian analyses: main analyses

*Reaction time*

**Table S5.** Bayesian model estimates from the reaction time model.

| | Estimate | Est. Error | l-95% CI | u-95% CI | R̂ | Bulk ESS | Tail ESS |
|---|---|---|---|---|---|---|---|
| Intercept | 356.85 | 3.42 | 350.14 | 363.63 | 1 | 1218 | 2047 |
| Right | -6.03 | 5.67 | -16.96 | 5.44 | 1 | 784 | 1176 |
| Left | 3.73 | 6.04 | -7.81 | 15.41 | 1 | 1121 | 1521 |
| Bilateral | -4.34 | 5.97 | -15.72 | 7.56 | 1.01 | 872 | 1940 |
| Epoch 1 | 20.79 | 0.44 | 19.93 | 21.67 | 1 | 5337 | 3456 |
| Epoch 2 | 8.05 | 0.45 | 7.17 | 8.95 | 1 | 6300 | 3678 |
| Epoch 3 | 0.66 | 0.45 | -0.2 | 1.57 | 1 | 5931 | 3437 |
| Epoch 4 | -3.92 | 0.44 | -4.77 | -3.05 | 1 | 5686 | 3512 |
| Epoch 5 | -7.97 | 0.46 | -8.86 | -7.08 | 1 | 5569 | 3519 |
| Triplet type | -4.46 | 0.2 | -4.86 | -4.07 | 1 | 6621 | 3438 |
| Right * Epoch 1 | 0.77 | 0.77 | -0.78 | 2.24 | 1 | 5073 | 3889 |
| Left * Epoch 1 | 3.13 | 0.8 | 1.55 | 4.69 | 1 | 5243 | 3726 |
| Bilateral * Epoch 1 | -1.25 | 0.8 | -2.78 | 0.31 | 1 | 5555 | 3624 |
| Right * Epoch 2 | -1.39 | 0.75 | -2.9 | 0.05 | 1 | 5206 | 4161 |
| Left * Epoch 2 | 1.11 | 0.8 | -0.43 | 2.65 | 1 | 5166 | 4001 |
| Bilateral * Epoch 2 | 0.23 | 0.8 | -1.34 | 1.81 | 1 | 4679 | 3305 |
| Right * Epoch 3 | -1.35 | 0.75 | -2.81 | 0.14 | 1 | 5414 | 3863 |
| Left * Epoch 3 | 0.62 | 0.79 | -0.94 | 2.16 | 1 | 5559 | 3743 |
| Bilateral * Epoch 3 | -0.63 | 0.78 | -2.14 | 0.91 | 1 | 4961 | 3671 |
| Right * Epoch 4 | -0.91 | 0.77 | -2.43 | 0.55 | 1 | 5862 | 3803 |
| Left * Epoch 4 | -3.05 | 0.8 | -4.58 | -1.46 | 1 | 5626 | 3618 |
| Bilateral * Epoch 4 | 2.08 | 0.78 | 0.59 | 3.58 | 1 | 5043 | 3739 |
| Right * Epoch 5 | 0.87 | 0.78 | -0.65 | 2.45 | 1 | 4613 | 2993 |
| Left * Epoch 5 | 0.77 | 0.83 | -0.82 | 2.42 | 1 | 4883 | 3588 |
| Bilateral * Epoch 5 | -0.27 | 0.79 | -1.83 | 1.31 | 1 | 5211 | 3202 |
| Right * Triplet type | 0.83 | 0.34 | 0.16 | 1.51 | 1 | 6932 | 3402 |
| Left * Triplet type | -0.78 | 0.36 | -1.46 | -0.09 | 1 | 6500 | 3318 |
| Bilateral * Triplet type | 0.48 | 0.36 | -0.23 | 1.18 | 1 | 6265 | 3190 |
| Epoch 1 * Triplet type | 2.8 | 0.45 | 1.93 | 3.66 | 1 | 6399 | 3573 |
| Epoch 2 * Triplet type | 1.56 | 0.46 | 0.66 | 2.47 | 1 | 6004 | 3771 |
| Epoch 3 * Triplet type | 0.75 | 0.45 | -0.12 | 1.64 | 1 | 5868 | 3777 |
| Epoch 4 * Triplet type | -1.25 | 0.46 | -2.14 | -0.34 | 1 | 4871 | 3846 |
| Epoch 5 * Triplet type | -2.82 | 0.46 | -3.73 | -1.91 | 1 | 6277 | 3305 |
| Right * Epoch 1 * Triplet type | 0.02 | 0.76 | -1.4 | 1.5 | 1 | 5542 | 3785 |
| Left * Epoch 1 * Triplet type | 0.44 | 0.8 | -1.09 | 2.02 | 1 | 5513 | 3377 |
| Bilateral * Epoch 1 * Triplet type | -0.04 | 0.78 | -1.6 | 1.45 | 1 | 5193 | 3703 |

| | | | | | | | |
|---|---|---|---|---|---|---|---|
| Right * Epoch 2 * Triplet type | -0.7 | 0.77 | -2.24 | 0.76 | 1 | 4385 | 3580 |
| Left * Epoch 2 * Triplet type | 0.15 | 0.79 | -1.43 | 1.7 | 1 | 4836 | 3191 |
| Bilateral * Epoch 2 * Triplet type | 0.13 | 0.79 | -1.46 | 1.69 | 1 | 4792 | 3642 |
| Right * Epoch 3 * Triplet type | -0.55 | 0.76 | -2.03 | 0.91 | 1 | 4954 | 3482 |
| Left * Epoch 3 * Triplet type | -0.39 | 0.79 | -1.95 | 1.15 | 1 | 5533 | 3904 |
| Bilateral * Epoch 3 * Triplet type | 0.46 | 0.78 | -1.09 | 2 | 1 | 4999 | 3840 |
| Right * Epoch 4 * Triplet type | 0.35 | 0.76 | -1.15 | 1.83 | 1 | 5242 | 3572 |
| Left * Epoch 4 * Triplet type | 0.45 | 0.8 | -1.08 | 1.99 | 1 | 5374 | 3867 |
| Bilateral * Epoch 4 * Triplet type | -1.09 | 0.78 | -2.64 | 0.47 | 1 | 5115 | 3518 |
| Right * Epoch 5 * Triplet type | 0.84 | 0.75 | -0.6 | 2.34 | 1 | 4518 | 3579 |
| Left * Epoch 5 * Triplet type | 0.46 | 0.81 | -1.11 | 2.05 | 1 | 4390 | 3432 |
| Bilateral * Epoch 5 * Triplet type | -0.09 | 0.79 | -1.65 | 1.47 | 1 | 5526 | 3797 |

**Notes.** Estimate = posterior mean; Est. Error = posterior standard deviation; CrI = 95% credible interval; R^ = potential scale reduction factor, ESS: effective sample size. For Group, the sham stimulation group served as the reference category; for Epoch, Epoch 6 served as the reference category.

**Table S6.** Bayesian post hoc pairwise comparison of the reaction time differences across epochs (Epoch main effect).

| contrast | | estimate | lower HPD | upper HPD |
|---|---|---|---|---|
| Epoch 1 | Epoch 2 | 12.75 | 11.47 | 14.12 |
| Epoch 1 | Epoch 3 | 20.14 | 18.77 | 21.45 |
| Epoch 1 | Epoch 4 | 24.72 | 23.42 | 2 6 |
| Epoch 1 | Epoch 5 | 28.7 6 | 27.3 6 | 30.08 |
| Epoch 1 | Epoch 6 | 38.42 | 37.02 | 39. 67 |
| Epoch 2 | Epoch 3 | 7.39 | 6.05 | 8.71 |
| Epoch 2 | Epoch 4 | 11.9 6 | 10. 63 | 13.35 |
| Epoch 2 | Epoch 5 | 1 6.02 | 14.58 | 17.33 |
| Epoch 2 | Epoch 6 | 25. 6 6 | 24.3 6 | 27.08 |
| Epoch 3 | Epoch 4 | 4.58 | 3.2 | 5.92 |
| Epoch 3 | Epoch 5 | 8. 62 | 7.2 6 | 10.07 |
| Epoch 3 | Epoch 6 | 18.27 | 1 6.92 | 19. 6 6 |
| Epoch 4 | Epoch 5 | 4.04 | 2.75 | 5.48 |
| Epoch 4 | Epoch 6 | 13.7 | 12.34 | 15.07 |
| Epoch 5 | Epoch 6 | 9. 64 | 8.2 6 | 10.9 6 |

**Note**. HPD = Highest Posterior Density, interval probability: 0.95

**Table S7.** Bayesian post hoc pairwise comparison of the reaction time differences across groups and epochs (Group * Epoch interaction).

| **Epoch 1** | | | | |
|---|---|---|---|---|
| **contrast** | | **estimate** | **lower HPD** | **upper HPD** |
| Sham | Right | -12.112 | -32.26 | 5.65 |
| Sham | Left | 0.374 | -17.36 | 19.31 |
| Sham | Bilateral | -9.466 | -27.41 | 10.15 |
| Right | Left | 12.542 | -6.8 | 31.86 |
| Right | Bilateral | 2.725 | -15.91 | 21.93 |
| Left | Bilateral | -9.77 | -28.6 | 9.44 |
| **Epoch 2** | | | | |
| **contrast** | | **estimate** | **lower HPD** | **upper HPD** |
| Sham | Right | -12.246 | -31.18 | 6.65 |
| Sham | Left | -3.235 | -21.86 | 14.74 |
| Sham | Bilateral | -14.26 | -31.64 | 5.46 |
| Right | Left | 9.001 | -10.73 | 27.8 |
| Right | Bilateral | -1.794 | -19.58 | 18.45 |
| Left | Bilateral | -10.832 | -29.57 | 8.84 |
| **Epoch 3** | | | | |
| **contrast** | | **estimate** | **lower HPD** | **upper HPD** |
| Sham | Right | -11.669 | -31.2 | 6.33 |
| Sham | Left | -2.351 | -21.6 | 15.35 |
| Sham | Bilateral | -15.74 | -33.94 | 3.9 |
| Right | Left | 9.393 | -9.64 | 29.4 |
| Right | Bilateral | -3.604 | -21.98 | 15.81 |
| Left | Bilateral | -13.109 | -32.8 | 5.68 |
| **Epoch 4** | | | | |
| **contrast** | | **estimate** | **lower HPD** | **upper HPD** |
| Sham | Right | -7.67 | -27.17 | 10.5 |
| Sham | Left | -4.648 | -23.21 | 13.65 |
| Sham | Bilateral | -15.544 | -34.16 | 3.58 |
| Right | Left | 3.162 | -16.21 | 22.46 |
| Right | Bilateral | -7.753 | -26.54 | 11.37 |
| Left | Bilateral | -10.872 | -31.15 | 7.1 |
| **Epoch 5** | | | | |
| **contrast** | | **estimate** | **lower HPD** | **upper HPD** |
| Sham | Right | -9.709 | -27.97 | 9.66 |
| Sham | Left | -0.493 | -18.81 | 18.1 |
| Sham | Bilateral | -10.476 | -28.4 | 9.18 |
| Right | Left | 9.223 | -10.02 | 28.89 |

| | | | | |
|---|---|---|---|---|
| Right | Bilateral | -0.83 | -19.16 | 18.36 |
| Left | Bilateral | -9.937 | -28.37 | 10.24 |

Epoch 6

| contrast | | estimate | lower HPD | upper HPD |
|---|---|---|---|---|
| Sham | Right | -5.052 | -24.59 | 13.05 |
| Sham | Left | 0.401 | -17.35 | 19.41 |
| Sham | Bilateral | -11.695 | -29.46 | 7.91 |
| Right | Left | 5.801 | -13.33 | 25.51 |
| Right | Bilateral | -6.289 | -24.59 | 13.38 |
| Left | Bilateral | -12.071 | -30.59 | 7.66 |

Notes. HPD = Highest Posterior Density, interval probability: 0.95

**Table S8.** Bayesian post hoc pairwise comparisons of reaction times groups (Group main effect).

| **contrast** | | **estimate** | **lower HPD** | **upper HPD** |
|---|---|---|---|---|
| Sham | Right | -9.8 | -30.2 | 7.14 |
| Sham | Left | -1.65 | -18.9 | 17.67 |
| Sham | Bilateral | -12.93 | -30.6 | 6.78 |
| Right | Left | 8.2 | -11 | 27.64 |
| Right | Bilateral | -2.93 | -21.5 | 16.09 |
| Left | Bilateral | -11.06 | -30.1 | 7.73 |

Notes. HPD = Highest Posterior Density, interval probability: 0.95

**Table S9.** Bayesian post hoc pairwise comparisons of reaction times for high- vs. low-probability triplets across epochs (Triplet type * Epoch interaction).

| | **H - L contrast estimate** | **lower HPD** | **upper HPD** |
|---|---|---|---|
| Epoch 1 | -3.31 | -5.17 | -1.43 |
| Epoch 2 | -5.81 | -7.68 | -3.87 |
| Epoch 3 | -7.41 | -9.31 | -5.51 |
| Epoch 4 | -11.39 | -13.4 | -9.5 |
| Epoch 5 | -14.54 | -16.56 | -12.56 |
| Epoch 6 | -10.99 | -12.78 | -8.93 |

Notes. H = high-probability triplet, L = low-probability triplet, HPD = Highest Posterior Density, interval probability: 0.95

**Table S10.** Bayesian post hoc pairwise comparisons of reaction times for high- vs. low-probability triplets across groups (Triplet type * Group interaction).

| **Group H-L contrast** | **estimate** | **lower HPD** | **upper HPD** |
|---|---|---|---|
| Sham-right | 3.207 | 1.146 | 5.43 |
| Sham-left | 0.705 | -1.645 | 2.853 |
| Sham-bilateral | 2.73 | 0.617 | 4.925 |
| Right-left | -2.523 | -4.752 | -0.134 |

| | | | |
|---|---|---|---|
| Right-bilateral | -0.496 | -2.615 | 1.863 |
| Left-bilateral | 2.002 | -0.242 | 4.27 |

Notes. HPD = Highest Posterior Density, interval probability: 0.95

**Table S11.** Bayesian post hoc pairwise comparisons of reaction times for high- vs. low-probability triplets across groups and epochs (Triplet type * Group * Epoch interaction).

| Epoch pairwise | Group pairwise | Triplet type pairwise | estimate | lower HPD | upper HPD |
|---|---|---|---|---|---|
| Epoch 1-Epoch 2 | Sham-right | H-L | 0.839 | -7 | 8.401 |
| Epoch 1-Epoch 3 | Sham-right | H-L | -0.474 | -7.79 | 7.23 |
| Epoch 1-Epoch 4 | Sham-right | H-L | -0.624 | -7.99 | 7.408 |
| Epoch 1-Epoch 5 | Sham-right | H-L | -1.561 | -8.83 | 6.828 |
| Epoch 1-Epoch 6 | Sham-right | H-L | -3.157 | -10.7 | 4.548 |
| Epoch 2-Epoch 3 | Sham-right | H-L | -1.347 | -9.35 | 6.111 |
| Epoch 2-Epoch 4 | Sham-right | H-L | -1.575 | -8.8 | 6.664 |
| Epoch 2-Epoch 5 | Sham-right | H-L | -2.498 | -10.5 | 4.8 |
| Epoch 2-Epoch 6 | Sham-right | H-L | -3.959 | -11.65 | 3.879 |
| Epoch 3-Epoch 4 | Sham-right | H-L | -0.143 | -7.62 | 7.741 |
| Epoch 3-Epoch 5 | Sham-right | H-L | -1.064 | -8.29 | 7.128 |
| Epoch 3-Epoch 6 | Sham-right | H-L | -2.706 | -9.91 | 5.324 |
| Epoch 4-Epoch 5 | Sham-right | H-L | -1.003 | -8.83 | 6.503 |
| Epoch 4-Epoch 6 | Sham-right | H-L | -2.596 | -10.39 | 4.872 |
| Epoch 5-Epoch 6 | Sham-right | H-L | -1.539 | -9.33 | 6.249 |
| Epoch 1-Epoch 2 | Sham-left | H-L | 1.741 | -5.82 | 9.606 |
| Epoch 1-Epoch 3 | Sham-left | H-L | 2.185 | -5.21 | 9.858 |
| Epoch 1-Epoch 4 | Sham-left | H-L | -2.738 | -10.37 | 4.662 |
| Epoch 1-Epoch 5 | Sham-left | H-L | -1.7 | -9.13 | 5.858 |
| Epoch 1-Epoch 6 | Sham-left | H-L | 1.253 | -6.58 | 8.727 |
| Epoch 2-Epoch 3 | Sham-left | H-L | 0.432 | -7.48 | 7.871 |
| Epoch 2-Epoch 4 | Sham-left | H-L | -4.528 | -11.97 | 3.06 |
| Epoch 2-Epoch 5 | Sham-left | H-L | -3.537 | -10.51 | 4.367 |
| Epoch 2-Epoch 6 | Sham-left | H-L | -0.538 | -8.41 | 7.082 |
| Epoch 3-Epoch 4 | Sham-left | H-L | -4.898 | -12.27 | 2.436 |
| Epoch 3-Epoch 5 | Sham-left | H-L | -3.842 | -11.87 | 3.436 |
| Epoch 3-Epoch 6 | Sham-left | H-L | -0.805 | -8.47 | 6.822 |
| Epoch 4-Epoch 5 | Sham-left | H-L | 1.021 | -7.2 | 8.1 |
| Epoch 4-Epoch 6 | Sham-left | H-L | 4.12 | -3.7 | 11.61 |
| Epoch 5-Epoch 6 | Sham-left | H-L | 3.079 | -4.66 | 10.663 |
| Epoch 1-Epoch 2 | Sham-bilateral | H-L | 3.157 | -4.37 | 10.823 |
| Epoch 1-Epoch 3 | Sham-bilateral | H-L | 3.023 | -4.53 | 10.743 |
| Epoch 1-Epoch 4 | Sham-bilateral | H-L | 0.801 | -6.59 | 8.406 |
| Epoch 1-Epoch 5 | Sham-bilateral | H-L | -3.214 | -10.87 | 3.954 |
| Epoch 1-Epoch 6 | Sham-bilateral | H-L | 1.576 | -5.8 | 9.362 |

| | | | | | |
|---|---|---|---|---|---|
| Epoch 2-Epoch 3 | Sham-bilateral | H-L | -0.116 | -7.85 | 7.59 |
| Epoch 2-Epoch 4 | Sham-bilateral | H-L | -2.324 | -9.76 | 5.52 |
| Epoch 2-Epoch 5 | Sham-bilateral | H-L | -6.352 | -13.83 | 0.727 |
| Epoch 2-Epoch 6 | Sham-bilateral | H-L | -1.402 | -9.14 | 6.25 |
| Epoch 3-Epoch 4 | Sham-bilateral | H-L | -2.225 | -9.64 | 5.509 |
| Epoch 3-Epoch 5 | Sham-bilateral | H-L | -6.12 | -13.33 | 1.602 |
| Epoch 3-Epoch 6 | Sham-bilateral | H-L | -1.293 | -9.2 | 6.546 |
| Epoch 4-Epoch 5 | Sham-bilateral | H-L | -4.008 | -11.65 | 3.07 |
| Epoch 4-Epoch 6 | Sham-bilateral | H-L | 0.969 | -6.86 | 8.161 |
| Epoch 5-Epoch 6 | Sham-bilateral | H-L | 4.833 | -3.24 | 12.115 |
| Epoch 1-Epoch 2 | Right-left | H-L | 0.871 | -6.95 | 8.92 |
| Epoch 1-Epoch 3 | Right-left | H-L | 2.689 | -5.29 | 10.224 |
| Epoch 1-Epoch 4 | Right-left | H-L | -2.115 | -10.24 | 5.724 |
| Epoch 1-Epoch 5 | Right-left | H-L | -0.172 | -8.11 | 7.566 |
| Epoch 1-Epoch 6 | Right-left | H-L | 4.474 | -3.66 | 12.167 |
| Epoch 2-Epoch 3 | Right-left | H-L | 1.689 | -6.08 | 9.65 |
| Epoch 2-Epoch 4 | Right-left | H-L | -3.075 | -10.76 | 4.824 |
| Epoch 2-Epoch 5 | Right-left | H-L | -1.031 | -8.71 | 6.902 |
| Epoch 2-Epoch 6 | Right-left | H-L | 3.554 | -4.62 | 11.06 |
| Epoch 3-Epoch 4 | Right-left | H-L | -4.719 | -12.5 | 2.923 |
| Epoch 3-Epoch 5 | Right-left | H-L | -2.828 | -11.16 | 5.01 |
| Epoch 3-Epoch 6 | Right-left | H-L | 1.779 | -6.03 | 9.643 |
| Epoch 4-Epoch 5 | Right-left | H-L | 1.934 | -6.09 | 9.517 |
| Epoch 4-Epoch 6 | Right-left | H-L | 6.645 | -0.86 | 14.314 |
| Epoch 5-Epoch 6 | Right-left | H-L | 4.687 | -3.57 | 12.228 |
| Epoch 1-Epoch 2 | Right-bilateral | H-L | 2.228 | -5.59 | 10.009 |
| Epoch 1-Epoch 3 | Right-bilateral | H-L | 3.472 | -4.49 | 11.063 |
| Epoch 1-Epoch 4 | Right-bilateral | H-L | 1.363 | -6.5 | 9.178 |
| Epoch 1-Epoch 5 | Right-bilateral | H-L | -1.638 | -8.98 | 6.43 |
| Epoch 1-Epoch 6 | Right-bilateral | H-L | 4.822 | -3.34 | 12.333 |
| Epoch 2-Epoch 3 | Right-bilateral | H-L | 1.207 | -6.4 | 9.39 |
| Epoch 2-Epoch 4 | Right-bilateral | H-L | -0.902 | -8.67 | 7.043 |
| Epoch 2-Epoch 5 | Right-bilateral | H-L | -3.874 | -11.63 | 3.692 |
| Epoch 2-Epoch 6 | Right-bilateral | H-L | 2.546 | -5.4 | 9.964 |
| Epoch 3-Epoch 4 | Right-bilateral | H-L | -2.136 | -10.02 | 5.874 |
| Epoch 3-Epoch 5 | Right-bilateral | H-L | -5.151 | -12.78 | 2.839 |
| Epoch 3-Epoch 6 | Right-bilateral | H-L | 1.399 | -6.66 | 9.09 |
| Epoch 4-Epoch 5 | Right-bilateral | H-L | -3.05 | -10.65 | 4.885 |
| Epoch 4-Epoch 6 | Right-bilateral | H-L | 3.422 | -4.41 | 11.232 |
| Epoch 5-Epoch 6 | Right-bilateral | H-L | 6.436 | -1.07 | 14.604 |
| Epoch 1-Epoch 2 | Left-bilateral | H-L | 1.435 | -6.41 | 8.997 |
| Epoch 1-Epoch 3 | Left-bilateral | H-L | 0.828 | -6.54 | 8.895 |

| | | | | | |
|---|---|---|---|---|---|
| Epoch 1-Epoch 4 | Left-bilateral | H-L | 3.497 | -4.11 | 11.306 |
| Epoch 1-Epoch 5 | Left-bilateral | H-L | -1.418 | -9.46 | 6.046 |
| Epoch 1-Epoch 6 | Left-bilateral | H-L | 0.409 | -7.36 | 7.902 |
| Epoch 2-Epoch 3 | Left-bilateral | H-L | -0.498 | -8.52 | 7.161 |
| Epoch 2-Epoch 4 | Left-bilateral | H-L | 2.123 | -5.39 | 10.076 |
| Epoch 2-Epoch 5 | Left-bilateral | H-L | -2.758 | -10.31 | 4.944 |
| Epoch 2-Epoch 6 | Left-bilateral | H-L | -1.074 | -8.96 | 6.696 |
| Epoch 3-Epoch 4 | Left-bilateral | H-L | 2.674 | -5.19 | 10.891 |
| Epoch 3-Epoch 5 | Left-bilateral | H-L | -2.297 | -10.61 | 4.873 |
| Epoch 3-Epoch 6 | Left-bilateral | H-L | -0.477 | -8.67 | 7.409 |
| Epoch 4-Epoch 5 | Left-bilateral | H-L | -5.031 | -12.54 | 2.785 |
| Epoch 4-Epoch 6 | Left-bilateral | H-L | -3.101 | -11.2 | 4.485 |
| Epoch 5-Epoch 6 | Left-bilateral | H-L | 1.835 | -6.22 | 9.444 |

**Note**. HPD = Highest Posterior Density, interval probability: 0.95

*Accuracy*

**Table S12.** Bayesian model estimates for the accuracy model.

| **Parameter** | **Estimate** | **Est. Error** | **95% Cri Lower** | **95% Cri Upper** | **$\hat{R}$** | **Bulk ESS** | **Tail ESS** |
|---|---|---|---|---|---|---|---|
| Intercept | 2.98 | 0.07 | 2.84 | 3.13 | 1.00 | 652 | 1381 |
| Right | -0.06 | 0.12 | -0.29 | 0.19 | 1.00 | 786 | 1509 |
| Left | 0.25 | 0.13 | 0.00 | 0.50 | 1.01 | 627 | 1188 |
| Bilateral | -0.21 | 0.13 | -0.48 | 0.04 | 1.01 | 471 | 782 |
| Epoch 1 | 0.23 | 0.03 | 0.18 | 0.28 | 1.00 | 4694 | 3634 |
| Epoch 2 | 0.08 | 0.02 | 0.03 | 0.13 | 1.00 | 5080 | 3523 |
| Epoch 3 | 0.02 | 0.02 | -0.03 | 0.07 | 1.00 | 5004 | 3434 |
| Epoch 4 | -0.05 | 0.02 | -0.10 | 0.00 | 1.00 | 5354 | 3579 |
| Epoch 5 | -0.09 | 0.02 | -0.14 | -0.05 | 1.00 | 5141 | 3659 |
| Triplet Type | 0.20 | 0.01 | 0.18 | 0.23 | 1.00 | 6356 | 3094 |
| Right * Epoch 1 | 0.05 | 0.04 | -0.03 | 0.13 | 1.00 | 4391 | 3302 |
| Left * Epoch 1 | -0.03 | 0.05 | -0.13 | 0.06 | 1.00 | 3707 | 3492 |
| Bilateral *Epoch 1 | -0.01 | 0.04 | -0.09 | 0.08 | 1.00 | 4031 | 3444 |
| Right * Epoch 2 | -0.06 | 0.04 | -0.13 | 0.02 | 1.00 | 4502 | 3781 |
| Left * Epoch 2 | 0.06 | 0.05 | -0.03 | 0.15 | 1.00 | 3773 | 3704 |

| | | | | | | | |
|---|---|---|---|---|---|---|---|
| Bilateral * Epoch 2 | 0.06 | 0.04 | -0.02 | 0.15 | 1.00 | 4151 | 3623 |
| Right * Epoch 3 | 0.03 | 0.04 | -0.05 | 0.11 | 1.00 | 4639 | 3718 |
| Left * Epoch 3 | 0.01 | 0.05 | -0.07 | 0.11 | 1.00 | 3897 | 3415 |
| Bilateral * Epoch 3 | -0.03 | 0.04 | -0.11 | 0.05 | 1.00 | 4274 | 3303 |
| Right * Epoch 4 | -0.03 | 0.04 | -0.10 | 0.05 | 1.00 | 4423 | 3527 |
| Left * Epoch 4 | 0.00 | 0.05 | -0.09 | 0.09 | 1.00 | 4187 | 3521 |
| Bilateral * Epoch 4 | 0.00 | 0.04 | -0.08 | 0.08 | 1.00 | 4459 | 3766 |
| Right * Epoch 5 | 0.00 | 0.04 | -0.07 | 0.07 | 1.00 | 4203 | 3584 |
| Left * Epoch 5 | 0.04 | 0.05 | -0.04 | 0.13 | 1.00 | 3813 | 2902 |
| Bilateral * Epoch 5 | -0.03 | 0.04 | -0.10 | 0.05 | 1.00 | 4232 | 3449 |
| Right * Triplet Type | 0.01 | 0.02 | -0.02 | 0.05 | 1.00 | 5152 | 3469 |
| Left * Triplet Type | 0.01 | 0.02 | -0.03 | 0.05 | 1.00 | 4704 | 3229 |
| Bilateral * Triplet Type | 0.00 | 0.02 | -0.03 | 0.04 | 1.00 | 4604 | 2972 |
| Epoch 1 * Triplet Type | -0.12 | 0.03 | -0.17 | -0.07 | 1.00 | 4514 | 3282 |
| Epoch 2 * Triplet Type | -0.02 | 0.02 | -0.07 | 0.03 | 1.00 | 5230 | 3548 |
| Epoch 3 * Triplet Type | -0.03 | 0.02 | -0.07 | 0.02 | 1.00 | 4885 | 3414 |
| Epoch 4 * Triplet Type | 0.04 | 0.02 | -0.01 | 0.08 | 1.00 | 5222 | 3635 |
| Epoch 5 * Triplet Type | 0.06 | 0.02 | 0.02 | 0.11 | 1.00 | 5479 | 3620 |
| Right * Epoch 1 * Triplet Type | -0.06 | 0.04 | -0.15 | 0.01 | 1.00 | 3504 | 3445 |
| Left * Epoch 1 * Triplet Type | 0.01 | 0.05 | -0.09 | 0.11 | 1.00 | 3516 | 3710 |
| Bilateral * Epoch 1 * Triplet Type | 0.03 | 0.04 | -0.05 | 0.11 | 1.00 | 3762 | 3488 |
| Right * Epoch 2 * Triplet Type | 0.03 | 0.04 | -0.05 | 0.10 | 1.00 | 3982 | 3496 |
| Left * Epoch 2 * Triplet Type | 0.06 | 0.05 | -0.03 | 0.16 | 1.00 | 3606 | 3447 |
| Bilateral * Epoch 2 * Triplet Type | -0.10 | 0.04 | -0.18 | -0.02 | 1.00 | 4223 | 3578 |

| | | | | | | | |
|---|---|---|---|---|---|---|---|
| Right * Epoch 3 * Triplet Type | 0.01 | 0.04 | -0.07 | 0.08 | 1.00 | 4308 | 3734 |
| Left * Epoch 3 * Triplet Type | 0.00 | 0.05 | -0.09 | 0.09 | 1.00 | 3878 | 3197 |
| Bilateral * Epoch 3 * Triplet Type | 0.06 | 0.04 | -0.02 | 0.13 | 1.00 | 4418 | 3706 |
| Right * Epoch 4 * Triplet Type | -0.01 | 0.04 | -0.09 | 0.06 | 1.00 | 3821 | 3402 |
| Left * Epoch 4 * Triplet Type | -0.01 | 0.04 | -0.10 | 0.07 | 1.00 | 4197 | 3770 |
| Bilateral * Epoch 4 * Triplet Type | -0.04 | 0.04 | -0.12 | 0.04 | 1.00 | 4651 | 3762 |
| Right * Epoch 5 * Triplet Type | 0.03 | 0.04 | -0.05 | 0.10 | 1.00 | 4054 | 2932 |
| Left * Epoch 5 * Triplet Type | -0.01 | 0.04 | -0.10 | 0.08 | 1.00 | 4080 | 3862 |
| Bilateral * Epoch 5 * Triplet Type | 0.01 | 0.04 | -0.06 | 0.08 | 1.00 | 4619 | 3773 |

**Note.** Estimate = posterior mean; Est. Error = posterior standard deviation; CrI = 95% credible interval; R^ = potential scale reduction factor, ESS: effective sample size. For Group, the sham stimulation group served as the reference category; for Epoch, Epoch 6 served as the reference category.

**Table S13.** Bayesian post hoc pairwise comparisons of accuracy for epochs (Epoch main effect).

| contrast | | estimate | lower HPD | upper HPD |
|---|---|---|---|---|
| Epoch 1 | Epoch 2 | 0.1518 | 0.07138 | 0.226 |
| Epoch 1 | Epoch 3 | 0.2104 | 0.13053 | 0.282 |
| Epoch 1 | Epoch 4 | 0.2782 | 0.1991 | 0.35 |
| Epoch 1 | Epoch 5 | 0.3203 | 0.24342 | 0.392 |
| Epoch 1 | Epoch 6 | 0.4116 | 0.3428 | 0.487 |
| Epoch 2 | Epoch 3 | 0.0594 | -0.01222 | 0.133 |
| Epoch 2 | Epoch 4 | 0.126 | 0.05132 | 0.198 |
| Epoch 2 | Epoch 5 | 0.1691 | 0.09521 | 0.237 |
| Epoch 2 | Epoch 6 | 0.2603 | 0.19037 | 0.332 |
| Epoch 3 | Epoch 4 | 0.068 | -0.00234 | 0.146 |
| Epoch 3 | Epoch 5 | 0.1109 | 0.04247 | 0.186 |
| Epoch 3 | Epoch 6 | 0.2011 | 0.13127 | 0.269 |
| Epoch 4 | Epoch 5 | 0.0423 | -0.02611 | 0.115 |
| Epoch 4 | Epoch 6 | 0.1339 | 0.06436 | 0.205 |
| Epoch 5 | Epoch 6 | 0.091 | 0.02913 | 0.164 |

Notes. HPD = Highest Posterior Density, interval probability: 0.95

**Table S14.** Bayesian post hoc pairwise comparisons of accuracy for groups (Group main effect).

| contrast | | estimate | lower HPD | upper HPD |
|---|---|---|---|---|
| Sham | Right | -0.309 | -0.6896 | 0.108 |
| Sham | Left | 0.151 | -0.2418 | 0.563 |
| Sham | Bilateral | -0.073 | -0.4816 | 0.31 |
| Right | Left | 0.454 | 0.0523 | 0.899 |
| Right | Bilateral | 0.233 | -0.1713 | 0.644 |
| Left | Bilateral | -0.229 | -0.6328 | 0.212 |
| Left | Bilateral | -0.229 | -0.6328 | 0.212 |

Notes. HPD = Highest Posterior Density, interval probability: 0.95

**Table S15.** Bayesian post hoc pairwise comparisons of accuracy for groups across epochs (Group * Epoch interaction).

| **Epoch 1** | | | | |
|---|---|---|---|---|
| **contrast** | | **estimate** | **lower HPD** | **upper HPD** |
| Sham | Right | -0.2251 | -0.6368 | 0.20873 |
| Sham | Left | 0.2159 | -0.1947 | 0.65212 |
| Sham | Bilateral | -0.0157 | -0.4388 | 0.4036 |
| Right | Left | 0.4309 | -0.0234 | 0.88861 |
| Right | Bilateral | 0.2039 | -0.2181 | 0.64097 |
| Left | Bilateral | -0.2318 | -0.6568 | 0.23783 |
| **Epoch 2** | | | | |
| **contrast** | | **estimate** | **lower HPD** | **upper HPD** |
| Sham | Right | -0.4232 | -0.8502 | 0.00123 |
| Sham | Left | 0.0312 | -0.4024 | 0.45014 |
| Sham | Bilateral | -0.0663 | -0.4803 | 0.34751 |
| Right | Left | 0.4519 | 0.0243 | 0.91518 |
| Right | Bilateral | 0.3596 | -0.0739 | 0.80021 |
| Left | Bilateral | -0.1006 | -0.5343 | 0.3528 |
| **Epoch 3** | | | | |
| **contrast** | | **estimate** | **lower HPD** | **upper HPD** |
| Sham | Right | -0.2888 | -0.6873 | 0.17472 |
| Sham | Left | 0.2136 | -0.2053 | 0.6348 |
| Sham | Bilateral | -0.0314 | -0.4302 | 0.40292 |
| Right | Left | 0.4971 | 0.065 | 0.95863 |
| Right | Bilateral | 0.2596 | -0.1852 | 0.68667 |
| Left | Bilateral | -0.2488 | -0.6867 | 0.19611 |
| **Epoch 4** | | | | |
| **contrast** | | **estimate** | **lower HPD** | **upper HPD** |
| Sham | Right | -0.3374 | -0.7525 | 0.09215 |
| Sham | Left | 0.1251 | -0.2782 | 0.5647 |
| Sham | Bilateral | -0.1269 | -0.5292 | 0.29412 |
| Right | Left | 0.455 | 0.0328 | 0.93737 |

| | | | | |
|---|---|---|---|---|
| Right | Bilateral | 0.2054 | -0.2504 | 0.61923 |
| Left | Bilateral | -0.2571 | -0.6865 | 0.18557 |

**Epoch 5**

| contrast | | estimate | lower HPD | upper HPD |
|---|---|---|---|---|
| Sham | Right | -0.3479 | -0.7276 | 0.10631 |
| Sham | Left | 0.1852 | -0.2285 | 0.59637 |
| Sham | Bilateral | -0.0541 | -0.4542 | 0.35729 |
| Right | Left | 0.5251 | 0.1154 | 0.99568 |
| Right | Bilateral | 0.2956 | -0.1228 | 0.73085 |
| Left | Bilateral | -0.2361 | -0.6995 | 0.18732 |

**Epoch 6**

| contrast | | estimate | lower HPD | upper HPD |
|---|---|---|---|---|
| Sham | Right | -0.2187 | -0.6379 | 0.19979 |
| Sham | Left | 0.1478 | -0.2568 | 0.56718 |
| Sham | Bilateral | -0.1489 | -0.5346 | 0.27825 |
| Right | Left | 0.3669 | -0.0446 | 0.82235 |
| Right | Bilateral | 0.0697 | -0.3566 | 0.50341 |
| Left | Bilateral | -0.2987 | -0.7338 | 0.13751 |
| Left | Bilateral | -0.2987 | -0.7338 | 0.13751 |

Notes. HPD = Highest Posterior Density, interval probability: 0.95

**Table S16.** Bayesian post hoc pairwise comparisons of accuracy for groups across triplet types (Group * Triplet type interaction).

| Group pairwise | Triplet type pairwise | estimate | lower HPD | upper HPD |
|---|---|---|---|---|
| Sham-right | H-L | 0.00183 | -0.1119 | 0.127 |
| Sham-left | H-L | 0.02875 | -0.0764 | 0.143 |
| Sham-bilateral | H-L | 0.08436 | -0.0234 | 0.197 |
| Right-left | H-L | 0.02676 | -0.0972 | 0.148 |
| Right-bilateral | H-L | 0.08073 | -0.0446 | 0.204 |
| Left-bilateral | H-L | 0.05528 | -0.0585 | 0.17 |

Notes. HPD = Highest Posterior Density, interval probability: 0.95

**Table S17.** Bayesian post hoc pairwise comparisons of accuracy for across epochs and triplet types (Epoch * Triplet type interaction).

| | H - L contrast estimate | lower HPD | upper HPD |
|---|---|---|---|
| Epoch 1 | 0.172 | 0.0597 | 0.286 |
| Epoch 2 | 0.369 | 0.2663 | 0.478 |
| Epoch 3 | 0.358 | 0.2547 | 0.461 |
| Epoch 4 | 0.487 | 0.3808 | 0.581 |
| Epoch 5 | 0.537 | 0.4317 | 0.634 |

| Epoch 6 | 0.531 | 0.4343 | 0.62 |
|---|---|---|---|

Notes. HPD = Highest Posterior Density, interval probability: 0.95

**Table S18.** Bayesian post hoc pairwise comparisons of accuracy for groups across epochs and triplet types (epoch * group * triplet type interaction).

| Epoch pairwise | Group pairwise | Triplet type pairwise | Estimate | lower HPD | upper HPD |
|---|---|---|---|---|---|
| Epoch 1-Epoch 2 | Sham-right | H-L | -0.07371 | -0.5378 | 0.3669 |
| Epoch 1-Epoch 3 | Sham-right | H-L | -0.15276 | -0.5889 | 0.3168 |
| Epoch 1-Epoch 4 | Sham-right | H-L | -0.14816 | -0.5699 | 0.3016 |
| Epoch 1-Epoch 5 | Sham-right | H-L | -0.22333 | -0.6404 | 0.2289 |
| Epoch 1-Epoch 6 | Sham-right | H-L | -0.2675 | -0.6768 | 0.1785 |
| Epoch 2-Epoch 3 | Sham-right | H-L | -0.07573 | -0.5043 | 0.345 |
| Epoch 2-Epoch 4 | Sham-right | H-L | -0.07498 | -0.4793 | 0.361 |
| Epoch 2-Epoch 5 | Sham-right | H-L | -0.15235 | -0.5845 | 0.2605 |
| Epoch 2-Epoch 6 | Sham-right | H-L | -0.19151 | -0.6281 | 0.2064 |
| Epoch 3-Epoch 4 | Sham-right | H-L | 0.00518 | -0.4143 | 0.4276 |
| Epoch 3-Epoch 5 | Sham-right | H-L | -0.07154 | -0.5039 | 0.3304 |
| Epoch 3-Epoch 6 | Sham-right | H-L | -0.11257 | -0.5064 | 0.3258 |
| Epoch 4-Epoch 5 | Sham-right | H-L | -0.07579 | -0.4623 | 0.3516 |
| Epoch 4-Epoch 6 | Sham-right | H-L | -0.11591 | -0.511 | 0.3053 |
| Epoch 5-Epoch 6 | Sham-right | H-L | -0.03695 | -0.4367 | 0.3799 |
| Epoch 1-Epoch 2 | Sham-left | H-L | -0.44541 | -0.8356 | -00328 |
| Epoch 1-Epoch 3 | Sham-left | H-L | -0.09239 | -0.4888 | 0.2952 |
| Epoch 1-Epoch 4 | Sham-left | H-L | -0.2452 | -0.6189 | 0.1581 |
| Epoch 1-Epoch 5 | Sham-left | H-L | -0.22679 | -0.5983 | 0.1685 |
| Epoch 1-Epoch 6 | Sham-left | H-L | -0.13499 | -0.4952 | 0.2663 |
| Epoch 2-Epoch 3 | Sham-left | H-L | 0.35594 | -0.0177 | 0.7515 |
| Epoch 2-Epoch 4 | Sham-left | H-L | 0.19651 | -0.1717 | 0.5847 |
| Epoch 2-Epoch 5 | Sham-left | H-L | 0.22237 | -0.151 | 0.5779 |
| Epoch 2-Epoch 6 | Sham-left | H-L | 0.31349 | -0.0621 | 0.6915 |
| Epoch 3-Epoch 4 | Sham-left | H-L | -0.15186 | -0.5522 | 0.2146 |
| Epoch 3-Epoch 5 | Sham-left | H-L | -0.13616 | -0.5092 | 0.2178 |
| Epoch 3-Epoch 6 | Sham-left | H-L | -0.04289 | -0.4005 | 0.3295 |
| Epoch 4-Epoch 5 | Sham-left | H-L | 0.0201 | -0.3192 | 0.4102 |
| Epoch 4-Epoch 6 | Sham-left | H-L | 0.10949 | -0.2388 | 0.4904 |
| Epoch 5-Epoch 6 | Sham-left | H-L | 0.09123 | -0.2564 | 0.4472 |
| Epoch 1-Epoch 2 | Sham-bilateral | H-L | -0.21488 | -0.6058 | 0.2221 |
| Epoch 1-Epoch 3 | Sham-bilateral | H-L | -0.31837 | -0.7403 | 0.076 |
| Epoch 1-Epoch 4 | Sham-bilateral | H-L | -0.03151 | -0.4405 | 0.3569 |
| Epoch 1-Epoch 5 | Sham-bilateral | H-L | -0.2875 | -0.6925 | 0.1053 |
| Epoch 1-Epoch 6 | Sham-bilateral | H-L | -0.24562 | -0.6672 | 0.1355 |

| | | | | | |
|---|---|---|---|---|---|
| Epoch 2-Epoch 3 | Sham-bilateral | H-L | -0.10646 | -0.4987 | 0.3046 |
| Epoch 2-Epoch 4 | Sham-bilateral | H-L | 0.18366 | -0.1949 | 0.5694 |
| Epoch 2-Epoch 5 | Sham-bilateral | H-L | -0.07556 | -0.4572 | 0.3051 |
| Epoch 2-Epoch 6 | Sham-bilateral | H-L | -0.03523 | -0.4102 | 0.3677 |
| Epoch 3-Epoch 4 | Sham-bilateral | H-L | 0.28643 | -0.1066 | 0.6695 |
| Epoch 3-Epoch 5 | Sham-bilateral | H-L | 0.02896 | -0.3466 | 0.4307 |
| Epoch 3-Epoch 6 | Sham-bilateral | H-L | 0.0678 | -0.3442 | 0.4396 |
| Epoch 4-Epoch 5 | Sham-bilateral | H-L | -0.25836 | -0.6417 | 0.1069 |
| Epoch 4-Epoch 6 | Sham-bilateral | H-L | -0.21785 | -0.5808 | 0.1724 |
| Epoch 5-Epoch 6 | Sham-bilateral | H-L | 0.04115 | -0.3058 | 0.4334 |
| Epoch 1-Epoch 2 | Right-left | H-L | -0.37497 | -0.8473 | 0.0845 |
| Epoch 1-Epoch 3 | Right-left | H-L | 0.06313 | -0.3884 | 0.5243 |
| Epoch 1-Epoch 4 | Right-left | H-L | -0.0952 | -0.5391 | 0.3518 |
| Epoch 1-Epoch 5 | Right-left | H-L | -0.00204 | -0.4417 | 0.4286 |
| Epoch 1-Epoch 6 | Right-left | H-L | 0.13421 | -0.302 | 0.5634 |
| Epoch 2-Epoch 3 | Right-left | H-L | 0.43814 | -0.0027 | 0.89 |
| Epoch 2-Epoch 4 | Right-left | H-L | 0.27811 | -0.1566 | 0.719 |
| Epoch 2-Epoch 5 | Right-left | H-L | 0.37123 | -0.0369 | 0.8283 |
| Epoch 2-Epoch 6 | Right-left | H-L | 0.5024 | 0.0587 | 0.9148 |
| Epoch 3-Epoch 4 | Right-left | H-L | -0.15995 | -0.5703 | 0.2886 |
| Epoch 3-Epoch 5 | Right-left | H-L | -0.06364 | -0.4656 | 0.3637 |
| Epoch 3-Epoch 6 | Right-left | H-L | 0.07115 | -0.342 | 0.475 |
| Epoch 4-Epoch 5 | Right-left | H-L | 0.09512 | -0.363 | 0.4687 |
| Epoch 4-Epoch 6 | Right-left | H-L | 0.23091 | -0.187 | 0.6177 |
| Epoch 5-Epoch 6 | Right-left | H-L | 0.13671 | -0.2562 | 0.5381 |
| Epoch 1-Epoch 2 | Right-bilateral | H-L | -0.13934 | -0.6008 | 0.3449 |
| Epoch 1-Epoch 3 | Right-bilateral | H-L | -0.16182 | -0.6336 | 0.3007 |
| Epoch 1-Epoch 4 | Right-bilateral | H-L | 0.12013 | -0.3286 | 0.5804 |
| Epoch 1-Epoch 5 | Right-bilateral | H-L | -0.06253 | -0.5504 | 0.379 |
| Epoch 1-Epoch 6 | Right-bilateral | H-L | 0.01532 | -0.4083 | 0.4814 |
| Epoch 2-Epoch 3 | Right-bilateral | H-L | -0.02687 | -0.4636 | 0.4247 |
| Epoch 2-Epoch 4 | Right-bilateral | H-L | 0.2516 | -0.1849 | 0.7149 |
| Epoch 2-Epoch 5 | Right-bilateral | H-L | 0.07251 | -0.3761 | 0.5156 |
| Epoch 2-Epoch 6 | Right-bilateral | H-L | 0.1523 | -0.29 | 0.5948 |
| Epoch 3-Epoch 4 | Right-bilateral | H-L | 0.28542 | -0.1418 | 0.7306 |
| Epoch 3-Epoch 5 | Right-bilateral | H-L | 0.10058 | -0.3362 | 0.5273 |
| Epoch 3-Epoch 6 | Right-bilateral | H-L | 0.18367 | -0.2158 | 0.6349 |
| Epoch 4-Epoch 5 | Right-bilateral | H-L | -0.18257 | -0.621 | 0.23 |
| Epoch 4-Epoch 6 | Right-bilateral | H-L | -0.09731 | -0.5081 | 0.3219 |
| Epoch 5-Epoch 6 | Right-bilateral | H-L | 0.0818 | -0.3219 | 0.5342 |
| Epoch 1-Epoch 2 | Left-bilateral | H-L | 0.23823 | -0.202 | 0.651 |
| Epoch 1-Epoch 3 | Left-bilateral | H-L | -0.22755 | -0.6361 | 0.1995 |
| Epoch 1-Epoch 4 | Left-bilateral | H-L | 0.21396 | -0.1666 | 0.6558 |

| | | | | | |
|---|---|---|---|---|---|
| Epoch 1-Epoch 5 | Left-bilateral | H-L | -0.06499 | -0.4547 | 0.3429 |
| Epoch 1-Epoch 6 | Left-bilateral | H-L | -0.11351 | -0.5216 | 0.305 |
| Epoch 2-Epoch 3 | Left-bilateral | H-L | -0.46482 | -0.8556 | -0.0656 |
| Epoch 2-Epoch 4 | Left-bilateral | H-L | -0.02144 | -0.4053 | 0.4096 |
| Epoch 2-Epoch 5 | Left-bilateral | H-L | -0.29896 | -0.6606 | 0.1165 |
| Epoch 2-Epoch 6 | Left-bilateral | H-L | -0.34563 | -0.742 | 0.0315 |
| Epoch 3-Epoch 4 | Left-bilateral | H-L | 0.44586 | 0.0498 | 0.8182 |
| Epoch 3-Epoch 5 | Left-bilateral | H-L | 0.17008 | -0.2089 | 0.5658 |
| Epoch 3-Epoch 6 | Left-bilateral | H-L | 0.11505 | -0.2833 | 0.489 |
| Epoch 4-Epoch 5 | Left-bilateral | H-L | -0.27624 | -0.6575 | 0.0869 |
| Epoch 4-Epoch 6 | Left-bilateral | H-L | -0.33065 | -0.6989 | 0.0634 |
| Epoch 5-Epoch 6 | Left-bilateral | H-L | -0.05063 | -0.449 | 0.3236 |

Notes. HPD = Highest Posterior Density, interval probability: 0.95

#### Reaction time variability results with data filtering

In the main analysis, we did not apply outlier filtering on the reaction time data, considering the large trial number and the relatively low number of potential outliers (with only erroneous trials excluded, 0.51% of the trials were above 700 ms). However, to rule out any suspicion that the results are driven by outliers, we re-ran the analysis with two different filtering methods: the more conservative Median Absolute Deviation (MAD) [1,2], and a fixed 1000 ms threshold [3].

*Median Absolute Deviation*

Using MAD to flag potential outlier trials followed the guidelines of [1], thus, the following formula: median(abs(values - median(sample))) / 0.6744897501960817. We flagged trials with values above 2.5 as outliers. This resulted in the highest reaction time in our sample to be 503 ms. This way, out of 184127 trials (all participants' all sequential epochs), 9839 (5.34%) were excluded. Of these 9839 trials, 2096 (21.3% of all excluded trials) belonged to the sham, 2602 (26.45%) belonged to the right, 1872 (19.03%) to the left, and 3269 (33.22%) to the bilaterally stimulated group.

With this filtering, the Levene test yielded a similar result as our main analysis on the unfiltered data ($F(3, 174284) = 142.28$, $p < 0.001$). Group differences also remained significant based on the pairwise post hoc test (Holm correction), the right and the bilateral DLPFC stimulated groups still showed significantly higher variances compared to the sham and the left DLPFC stimulated groups ($p_{Holm} < 0.001$ in each case). See Figure S3 panel A for details.

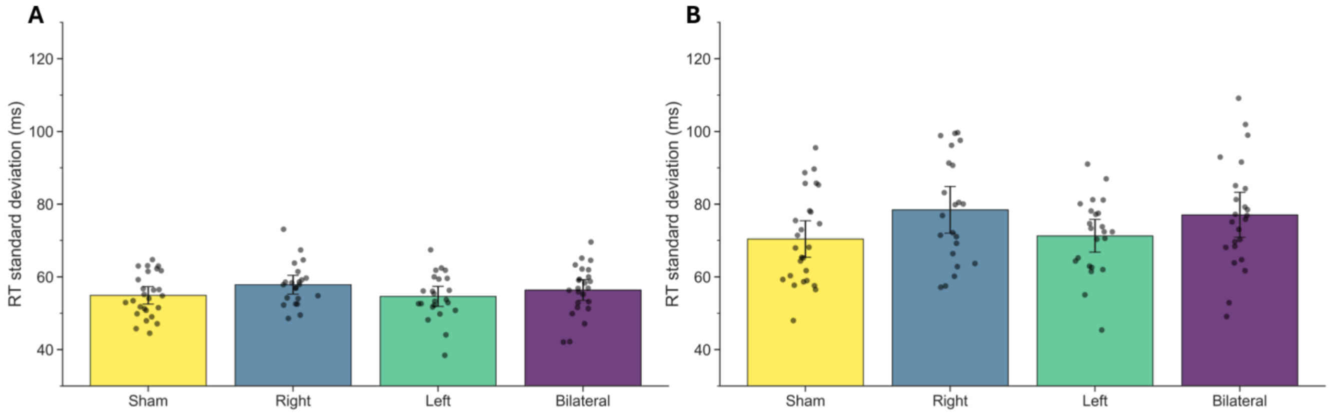


**Figure S3.** RT standard deviation with MAD filtering (panel A) and 1000 ms RT filtering (panel B). On both figures, the y-axis represents RT standard deviation, the x-axis and the colors depict the four groups. Black dots represent individual RT standard deviation data.

*Fixed threshold filtering*

Excluding trials with RTs above 1000 ms resulted in the loss of 124 trials out of 184127 (0.07%). Of these 124 trials, 28 (22.58% of all excluded trials) belonged to the sham, 43 (34.68%) belonged to the right, 16 (12.9%) to the left, and 37 (29.84%) to the bilaterally stimulated group. Performing the Levene test on the data filtered this way also supported the previous results ($F(3, 183998) = 286.76$, $p < 0.001$, post hoc pairwise comparisons $p_{Holm} < 0.001$).
In both cases, results matched with those yielded by the Brown–Forsythe test we additionally ran. Taken all of these results together, data filtering did not change the results reported in the main manuscript. See Figure S3 panel A for details.

**Speed-accuracy trade-off**

To test whether the groups differed in RT or accuracy, we ran some additional analyses. First, before the first stimulation, all participants completed a familiarization phase consisting of three randomized ASRT blocks without an embedded sequence. These blocks provide a genuine pre-stimulation measurement of general response speed and accuracy, although by construction they cannot index statistical learning. To make sure that our groups do not differ in the baseline RT and accuracy measured during familiarization, we ran two models: a linear mixed-effect model for RT, and a generalized linear mixed-effect model for accuracy. Group was the fixed factor, and participant number the random factor in both (random intercept included), while the outcome variables were the trial-wise RT or accuracy. We had to exclude one participant from these analysis, as their file was compromised. We found no significant group differences neither in the RT ($F(3,90.03) = 0.87$, $p = 0.458$), nor in the accuracy($\chi^2(3) = 5.32$, $p = 0.150$) during this familiarization phase, indicating that groups were comparable during the familiarization phase.

To further exclude the option that a group difference in the speed-accuracy trade-off (SATO) would be present in our data, we computed a score to quantify to which extent participants favoured accuracy or RT based on [4]. First, we calculated the z-scores of the median RT by epoch and the mean error rate, then we subtracted the latter from the former ($SATO = RT_z - error_z$). Positive scores mean a performance focused more on accuracy, negative scores indicate focus on speed, values near zero show a balanced performance.

We ran a frequentist and a Bayesian linear mixed model (*brm()* function of the *brms* package in R) using this value as the outcome, Group and Epoch as fixed factors, and Participant number as the random factor (the model including the random intercept). For the Bayesian model, we applied 3 chains with 4000 iterations and 500 warm ups to reach acceptable effective sample sizes, and the *brms* default priors. We found that the Epoch main effect was significant ($F(5, 455) = 90.01$, $p < 0.001$) due to a gradual decrease of the SATO score across Epoch 1-4 (post hoc $p_{Holm} = 0.047$), indicating an increasingly speed-focused performance. However, neither the Group main effect ($F(3, 91) = 1.45$, $p = 0.233$), nor the Group * Epoch interaction ($F(15, 455) = 0.71$, $p = 0.772$) was significant. The Bayesian results reflected a similar pattern, see Supplementary Tables S19-S22. See Figure S4 for details. These results make it unlikely that any of our main claims would be undermined by different speed-accuracy trade-off strategies across groups.

**Table S19.** Bayesian model estimates from the SATO model.

| | **Estimate** | **Estimate Error** | **l-95% CI** | **u-95% CI** | **$\hat{R}$** | **Bulk ESS** | **Tail ESS** |
|---|---|---|---|---|---|---|---|
| Intercept | 0 | 0.15 | -0.29 | 0.29 | 1 | 454 | 933 |
| Right | -0.28 | 0.26 | -0.8 | 0.23 | 1.01 | 362 | 721 |
| Left | 0.45 | 0.28 | -0.11 | 1 | 1.01 | 535 | 1179 |
| Bilateral | -0.32 | 0.29 | -0.87 | 0.25 | 1.01 | 385 | 583 |
| Epoch 1 | 0.89 | 0.05 | 0.79 | 1 | 1 | 5795 | 7083 |
| Epoch 2 | 0.35 | 0.06 | 0.24 | 0.46 | 1 | 6611 | 6942 |
| Epoch 3 | 0.04 | 0.05 | -0.07 | 0.15 | 1 | 6070 | 7155 |
| Epoch 4 | -0.2 | 0.05 | -0.31 | -0.1 | 1 | 6554 | 6786 |
| Epoch 5 | -0.37 | 0.05 | -0.48 | -0.26 | 1 | 6178 | 6856 |
| Right*Epoch 1 | 0.08 | 0.09 | -0.1 | 0.27 | 1 | 4710 | 5742 |
| Left*Epoch 1 | -0.04 | 0.1 | -0.24 | 0.15 | 1 | 5198 | 7131 |
| Bilateral*Epoch 1 | 0.04 | 0.1 | -0.14 | 0.23 | 1 | 5362 | 6020 |
| Right*Epoch 2 | -0.11 | 0.09 | -0.29 | 0.08 | 1 | 5464 | 7270 |
| Left*Epoch 2 | 0.11 | 0.1 | -0.08 | 0.29 | 1 | 5492 | 6377 |
| Bilateral*Epoch 2 | 0.07 | 0.1 | -0.11 | 0.26 | 1 | 5822 | 7066 |
| Right*Epoch 3 | 0.03 | 0.09 | -0.15 | 0.21 | 1 | 4965 | 6657 |
| Left*Epoch 3 | 0.02 | 0.1 | -0.17 | 0.21 | 1 | 4869 | 6272 |
| Bilateral*Epoch 3 | -0.05 | 0.1 | -0.24 | 0.14 | 1 | 4768 | 6646 |
| Right*Epoch 4 | -0.08 | 0.09 | -0.26 | 0.1 | 1 | 5480 | 6182 |
| Left*Epoch 4 | -0.06 | 0.1 | -0.25 | 0.13 | 1 | 5254 | 6550 |
| Bilateral*Epoch 4 | 0.02 | 0.1 | -0.16 | 0.21 | 1 | 4395 | 5997 |
| Right*Epoch 5 | 0.06 | 0.09 | -0.12 | 0.24 | 1 | 4614 | 6058 |
| Left*Epoch 5 | 0.13 | 0.1 | -0.06 | 0.33 | 1 | 4730 | 6142 |
| Bilateral*Epoch 5 | -0.1 | 0.1 | -0.29 | 0.09 | 1 | 4385 | 5883 |

**Note.** Estimate = posterior mean; Est. Error = posterior standard deviation; CrI = 95% credible interval; R^ = potential scale reduction factor, ESS: effective sample size. For Group, the sham stimulation group served as the reference category; for Epoch, Epoch 6 served as the reference category.

**Table S20.** Epoch main effect post hoc analysis for the SATO Bayesian model

| contrast | | estimate | lower HPD | upper HPD |
|---|---|---|---|---|
| Epoch 1 | Epoch 2 | 0.543 | 0.37245 | 0.706 |
| Epoch 1 | Epoch 3 | 0.852 | 0.69226 | 1.022 |
| Epoch 1 | Epoch 4 | 1.094 | 0.93431 | 1.263 |
| Epoch 1 | Epoch 5 | 1.258 | 1.08963 | 1.421 |
| Epoch 1 | Epoch 6 | 1.597 | 1.44165 | 1.77 |
| Epoch 2 | Epoch 3 | 0.309 | 0.13759 | 0.474 |
| Epoch 2 | Epoch 4 | 0.551 | 0.3874 | 0.716 |
| Epoch 2 | Epoch 5 | 0.717 | 0.55052 | 0.889 |
| Epoch 2 | Epoch 6 | 1.055 | 0.88967 | 1.221 |
| Epoch 3 | Epoch 4 | 0.242 | 0.0761 | 0.405 |
| Epoch 3 | Epoch 5 | 0.407 | 0.24632 | 0.571 |
| Epoch 3 | Epoch 6 | 0.745 | 0.57467 | 0.904 |
| Epoch 4 | Epoch 5 | 0.165 | 0.00643 | 0.332 |
| Epoch 4 | Epoch 6 | 0.503 | 0.34115 | 0.666 |
| Epoch 5 | Epoch 6 | 0.339 | 0.1754 | 0.498 |

**Note**. HPD = Highest Posterior Density, interval probability: 0.95

**Table S21.** Group main effect post hoc analysis for the SATO Bayesian model

| contrast | | estimate | lower HPD | upper HPD |
|---|---|---|---|---|
| Sham | Right | -0.731 | -1.58 | 0.14 |
| Sham | Left | 0.0398 | -0.822 | 0.888 |
| Sham | Bilateral | -0.4341 | -1.272 | 0.426 |
| Right | Left | 0.7748 | -0.126 | 1.704 |
| Right | Bilateral | 0.2974 | -0.571 | 1.226 |
| Left | Bilateral | -0.4937 | -1.356 | 0.434 |

**Note**. HPD = Highest Posterior Density, interval probability: 0.95

**Table S22.** Epoch*group interaction post hoc analysis for the SATO Bayesian model

| **Epoch 1** | | | | |
|---|---|---|---|---|
| **contrast** | | **estimate** | **lower HPD** | **upper HPD** |
| Sham | Right | -0.603 | -1.5526 | 0.2764 |
| Sham | Left | 0.0803 | -0.82116 | 1.0032 |
| Sham | Bilateral | -0.2698 | -1.16998 | 0.6118 |
| Right | Left | 0.6842 | -0.27147 | 1.6476 |
| Right | Bilateral | 0.3366 | -0.60796 | 1.2857 |
| Left | Bilateral | -0.3624 | -1.2506 | 0.659 |
| **Epoch 2** | | | | |

| contrast | | estimate | lower HPD | upper HPD |
|---|---|---|---|---|
| Sham | Right | -0.9386 | -1.84235 | -0.0277 |
| Sham | Left | -0.1379 | -1.08901 | 0.7474 |
| Sham | Bilateral | -0.4605 | -1.39727 | 0.3773 |
| Right | Left | 0.8111 | -0.19586 | 1.745 |
| Right | Bilateral | 0.4764 | -0.50432 | 1.3944 |
| Left | Bilateral | -0.3446 | -1.26463 | 0.6277 |
| **Epoch 3** | | | | |
| **contrast** | | **estimate** | **lower HPD** | **upper HPD** |
| Sham | Right | -0.7267 | -1.65227 | 0.2027 |
| Sham | Left | 0.1226 | -0.79288 | 1.0134 |
| Sham | Bilateral | -0.411 | -1.31848 | 0.499 |
| Right | Left | 0.8382 | -0.18616 | 1.7658 |
| Right | Bilateral | 0.3101 | -0.67774 | 1.2051 |
| Left | Bilateral | -0.5432 | -1.48729 | 0.418 |
| **Epoch 4** | | | | |
| **contrast** | | **estimate** | **lower HPD** | **upper HPD** |
| Sham | Right | -0.7488 | -1.62965 | 0.1978 |
| Sham | Left | -0.061 | -0.95776 | 0.8737 |
| Sham | Bilateral | -0.6247 | -1.52719 | 0.2713 |
| Right | Left | 0.6861 | -0.29511 | 1.6403 |
| Right | Bilateral | 0.1178 | -0.83551 | 1.048 |
| Left | Bilateral | -0.58 | -1.51257 | 0.3886 |
| **Epoch 5** | | | | |
| **contrast** | | **estimate** | **lower HPD** | **upper HPD** |
| Sham | Right | -0.8091 | -1.69314 | 0.1148 |
| Sham | Left | 0.197 | -0.68797 | 1.1297 |
| Sham | Bilateral | -0.2846 | -1.25577 | 0.5203 |
| Right | Left | 1.007 | -0.00646 | 1.9253 |
| Right | Bilateral | 0.5198 | -0.40966 | 1.4502 |
| Left | Bilateral | -0.5013 | -1.41253 | 0.4595 |
| **Epoch 6** | | | | |
| **contrast** | | **estimate** | **lower HPD** | **upper HPD** |
| Sham | Right | -0.57 | -1.48054 | 0.3583 |
| Sham | Left | 0.0366 | -0.84599 | 0.98 |
| Sham | Bilateral | -0.5424 | -1.4456 | 0.354 |
| Right | Left | 0.6071 | -0.35269 | 1.5908 |
| Right | Bilateral | 0.0214 | -0.89839 | 1.0047 |
| Left | Bilateral | -0.5961 | -1.5704 | 0.326 |

**Note**. HPD = Highest Posterior Density,  interval probability: 0.95

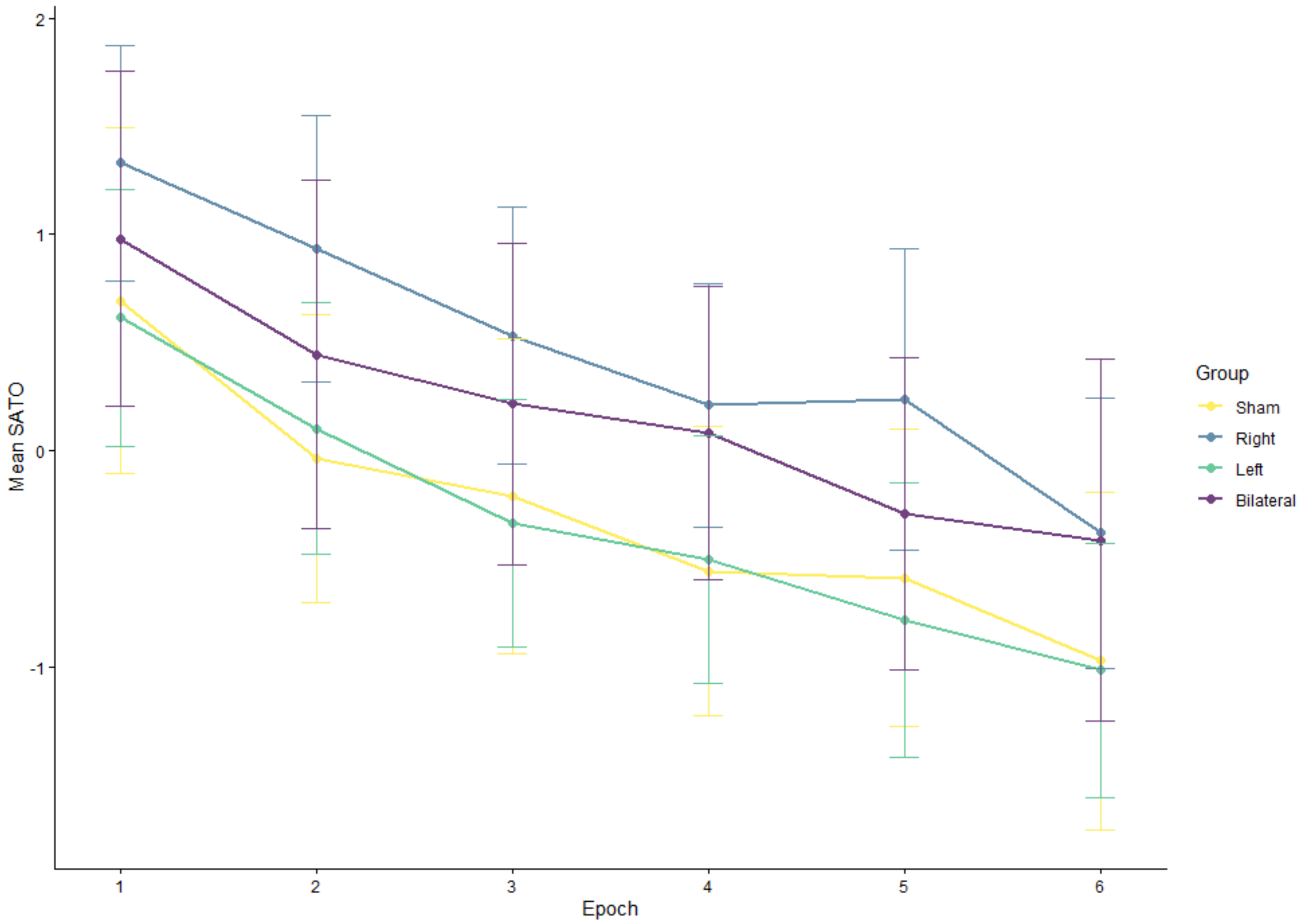


**Figure S4.** Speed-accuracy trade-off value in the four groups across epochs. Speed–accuracy trade-off (SATO) values across six epochs in the four stimulation groups. Lines represent mean SATO values for the sham (yellow), right (blue), left (green), and bilateral (purple) DLPFC stimulation groups. Error bars represent 95% confidence intervals.

#### References


Hann, F., Pesthy, O., Brezóczki, B., Vékony, T., Nagy, C. A., Sapey-Triomphe, L. A., Tóth-Fáber, E., Farkas, B. C., Farkas, K., & Németh, D. (2025). Autistic traits relate to speed/accuracy trade-off but not statistical learning and updating. *Scientific Reports 2025 15:1*, *15*(1), 32001-. https://doi.org/10.1038/s41598-025-16138-7

Krajcsi, A. (2021). *CogStat - An automatic analysis statistical software*. https://doi.org/10.31234/OSF.IO/HNMSQ

Leys, C., Ley, C., Klein, O., Bernard, P., & Licata, L. (2013). Detecting outliers: Do not use standard deviation around the mean, use absolute deviation around the median. *Journal of Experimental Social Psychology*, *49*(4), 764–766. https://doi.org/10.1016/J.JESP.2013.03.013

Zolnai, T., David, D. R., Pesthy, O., Nemeth, M., Kiss, M., Nagy, M., & Nemeth, D. (2022). Measuring statistical learning by eye-tracking. *Experimental Results*, *3*, e10. https://doi.org/10.1017/EXP.2022.8
